# A ubiquitin-based mechanism for the oligogenic inheritance of heterotaxy and heart defects

**DOI:** 10.1101/2020.05.25.113944

**Authors:** Jennifer H. Kong, Cullen B. Young, Ganesh V. Pusapati, Chandni B. Patel, Sebastian Ho, Arunkumar Krishnan, Jiuann-Huey Ivy Lin, William Devine, Anne Moreau de Bellaing, Tejas S. Athni, L. Aravind, Teresa M. Gunn, Cecilia W. Lo, Rajat Rohatgi

**Affiliations:** Departments of Biochemistry and Medicine, Stanford University School of Medicine, Stanford, California, United States of America; Department of Developmental Biology, University of Pittsburgh School of Medicine, Pittsburgh, Pennsylvania, United States of America; Department of Pediatric Cardiology, Necker-Sick Children Hospital and The University of Paris Descartes, Paris, France; McLaughlin Research Institute, Great Falls, Montana, United States of America; National Center for Biotechnology Information, National Library of Medicine, National Institutes of Health, Bethesda, MD 20894, USA

## Abstract

The etiology of congenital heart defects (CHDs), amongst the most common human birth defects, is poorly understood partly because of its complex genetic architecture. Here we show that two genes previously implicated in CHDs, *Megf8* and *Mgrn1*, interact genetically and biochemically to regulate the strength of Hedgehog signaling in target cells. MEGF8, a single-pass transmembrane protein, and MGRN1, a RING superfamily E3 ligase, assemble to form a transmembrane ubiquitin ligase complex that catalyzes the ubiquitination and degradation of the Hedgehog pathway transducer Smoothened. Homozygous *Megf8* and *Mgrn1* mutations increased Smoothened abundance and elevated sensitivity to Hedgehog ligands. While mice heterozygous for loss-of-function *Megf8* or *Mgrn1* mutations were normal, double heterozygous embryos exhibited an incompletely penetrant syndrome of CHDs with heterotaxy. Thus, genetic interactions between components of a receptor-like ubiquitin ligase complex that tunes morphogen signaling strength can cause a birth defect syndrome inherited in an oligogenic pattern.

## Introduction

Morphogens are secreted ligands that influence developmental outcomes (differentiation, patterning or morphogenesis) in a dose-dependent manner. For example, temporal and spatial gradients of Hedgehog (Hh) ligands (like Sonic Hedgehog, SHH) pattern the spinal cord and limb during development. Varying concentrations or durations of morphogen exposure produce different cellular outcomes by changing the strength or persistence of signaling in target cells (Harfe et al., 2004; Stamataki et al., 2005). Much of the focus in morphogen signaling has been on understanding how ligands like SHH are produced and distributed across tissues to form gradients. However, signaling strength in target cells is a function of both ligand exposure and ligand sensitivity. Much less is known about mechanisms that function in target cells to modulate ligand reception and whether such mechanisms are damaged in developmental diseases.

In recent CRISPR screens for regulators of Hh signaling, we discovered several proteins that function in target cells to attenuate signaling strength. Because of similarities in their loss-of-function phenotypes, we focus here on three of these proteins: Multiple Epidermal Growth Factor-like Domains 8 (MEGF8), a type I single-pass transmembrane protein, and two paralogous RING superfamily E3 ubiquitin ligases, Mahogunin Ring Finger 1 (MGRN1) and RNF157. *Megf8* was identified as a regulator of both left-right patterning and cardiac morphogenesis in mouse genetic screens (Aune et al., 2008; Li et al., 2015; Zhang et al., 2009). Human mutations in *MEGF8* result in Carpenter syndrome, an autosomal recessive syndrome similarly characterized by heterotaxy (defects in left-right patterning), severe congenital heart defects (CHDs), preaxial digit duplication, and skeletal defects (Twigg et al., 2012). Unlike many other genes associated with heterotaxy, loss of *Megf8* does not result in any detectable defects in either primary or motile cilia (Aune et al., 2008; Pusapati et al., 2018a; Zhang et al., 2009).

Loss of MGRN1 was also previously shown to cause CHDs and heterotaxy with partial penetrance in mice (Cota et al., 2006). How MEGF8 and MGRN1 regulate these critical developmental events has remained unknown for over a decade. We investigated the biochemical and biological functions of MEGF8, MGRN1 and RNF157 using a combination of mechanistic studies in cultured cells and mouse genetics. MEGF8, MGRN1 and RNF157 anchor a novel ubiquitination pathway that regulates the sensitivity of target cells to Hh ligands. They assemble into an unusual transmembrane E3 ubiquitin ligase complex that functions as a traffic control system for signaling receptors, including the Hh transducer Smoothened (SMO). Mouse studies revealed striking genetic interactions and gene dosage effects involving *Megf8*, *Mgrn1* and *Rnf157* that impact the penetrance of a wide spectrum of birth defects, including congenital heart defects (CHDs), heterotaxy, skeletal defects, and limb anomalies. Our work shows how genetic interactions can arise from biochemical mechanisms that regulate signaling strength, a conclusion with broader implications for understanding the molecular pathophysiology of human diseases in which oligogenic interactions are emerging as an important mechanism of heritability.

## Results

### *Megf8* and *Mgrn1* are negative regulators of Hedgehog signaling

Amongst the top gene hits identified in our genome-wide screen for attenuators of Hh signaling (Pusapati et al., 2018a), we pursued a detailed analysis of *Megf8* and *Mgrn1* because of similarities in their loss of function phenotypes (**Fig. S1A**). In both NIH/3T3 fibroblasts and cultured neural progenitor cells (NPCs), CRISPR-mediated loss-of-function mutations in *Megf8* and *Mgrn1* resulted in an elevated response to Sonic hedgehog (SHH) ligands caused by the accumulation of SMO at the cell surface and primary cilium (Pusapati et al., 2018a). To determine if MEGF8 and MGRN1 can attenuate Hh signaling in a more physiological context, we isolated primary mouse embryonic fibroblasts (pMEFs) from embryos homozygous for previously characterized mutant alleles of *Megf8* (C193R) or *Mgrn1* (md-nc) (**Fig. S1A**) (He et al., 2003; Phillips, 1963; Zhang et al., 2009). As we observed in NIH/3T3 cells, *Megf8^C193R/C193R^* and *Mgrn1^md-nc^*^/^*^md-nc^* (hereafter referred to as *Megf8^m/m^* and *Mgrn1^m^*^/^*^m^*) pMEFs were more sensitive to SHH. When exposed to a sub-saturating concentration of SHH (1 nM), *Gli1* (a direct Hh target gene) was only partially induced in wild-type pMEFs, but this same low concentration induced *Gli1* to maximum levels in *Megf8^m/m^* and *Mgrn1^m/m^* pMEFs (**Figs. S1B and S1C**).

Heightened SHH sensitivity was likely caused by an elevated abundance of SMO in the primary cilia of *Megf8^m/m^* and *Mgrn1^m/m^* pMEFs, both in the absence of SHH and when signaling was activated by pathway agonists (**Figs. S1D and S1E**).

The accumulation of ectopic ciliary SMO was also observed in multiple tissues within *Megf8^m/m^* and *Mgrn1^m/m^* embryos (**Fig. S2**). In wild-type embryos, Hh signaling activity is restricted to early embryonic development and by e12.5 is turned off in most tissues, resulting in ciliary SMO restricted to cells that were exposed to only the highest concentrations of SHH, like the progenitor cells within the ventral neural tube (**Fig. S2**) (Corbit et al., 2005; Rohatgi et al., 2007a). In contrast, SMO was concentrated in the primary cilia of nearly all *Megf8^m/m^* embryonic tissues, regardless of whether it had been exposed to Hh ligands (**Fig. S2**). Tissues from *Mgrn1^m/m^* embryos did not have widespread accumulation of ciliary SMO (**Fig. S2**). However, ciliary SMO was inappropriately present sporadically in the dorsal neural tube and brain of *Mgrn1^m/m^* embryos (**Fig. S2**), consistent with our observation that *Mgrn1^-/-^* NPCs exhibited a moderately elevated response to SHH (Pusapati et al., 2018a). The difference in the severity of the Hh signaling phenotype between *Megf8^m/m^* and *Mgrn1^m/m^* pMEFs and embryos (**Figs. S1 and S2**) is consistent with our previous results comparing *Mgrn1*^-/-^ and *Megf8*^-/-^ NIH/3T3 cells (Pusapati et al., 2018a). As reported previously, the loss of MEGF8 or MGRN1 did not cause any defects in the number or the structure of primary cilia (Pusapati et al., 2018b; Zhang et al., 2009).

### *Rnf157* is a genetic modifier of *Mgrn1*

In both mice and cultured cells, loss of MEGF8 consistently produced stronger phenotypes than the loss of MGRN1 (**Figs. S1 and S2**), suggesting the involvement of additional genes. The reported penetrance and expressivity of CHDs, heterotaxy, and preaxial digit duplication was much higher in *Megf8^m/m^* embryos compared to *Mgrn1^m/m^* embryos (Cota et al., 2006; Zhang et al., 2009) (**Table S1**). Similarly in NIH/3T3 cells, when compared to the loss of MGRN1, the loss of MEGF8 resulted in more Hh signaling activity at baseline and a greater abundance of SMO at the plasma and ciliary membranes (**Figs. 1A and 1C**). Evolutionary sequence analysis indicated that RNF157, which also encodes a RING superfamily E3 ligase, is a vertebrate-specific paralog of MGRN1 (**Fig. 1B**) (Pusapati et al., 2018a). Although MGRN1 is more widely distributed, found amongst almost all major eukaryotic lineages, MGRN1 and RNF157 share a RING domain and a distinctive predicted substrate-binding domain that is unique amongst other members of the RING superfamily (**Figs. 1B and S3A**).

**Figure 1:**
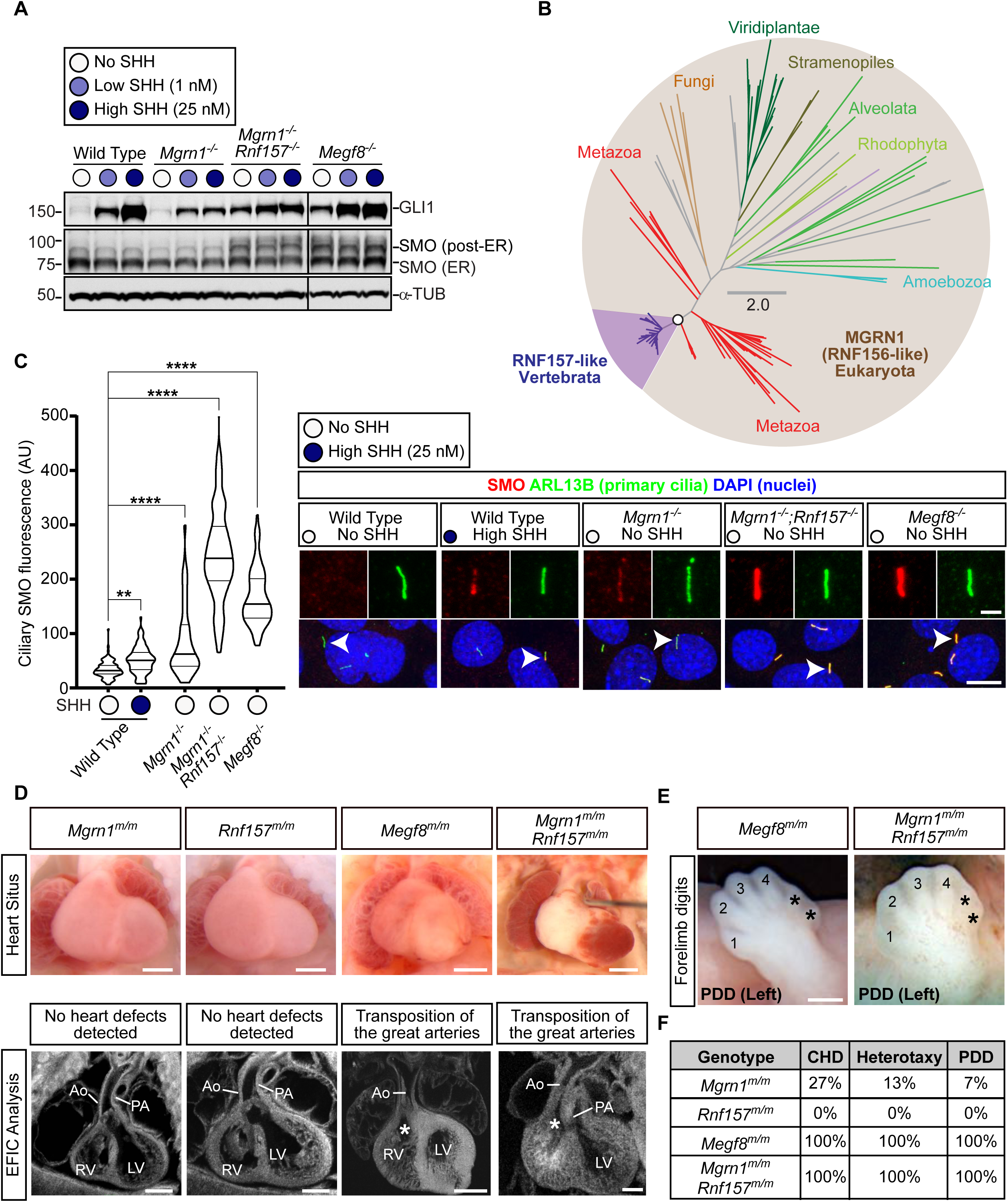
RNF157 partially compensates for the loss of MGRN1. **(A)** Immunoblots of wild-type, *Mgrn1*^-/-^, *Mgrn1*^-/-^;*Rnf157*^-/-^, and *Megf8*^-/-^ NIH/3T3 cells treated with no SHH (0 nM), low SHH (1 nM), or high SHH (25 nM). Immunoblots show GLI1 (a metric for signaling strength), SMO, and α-Tubulin (α-TUB) as a loading control. The fraction of SMO that has traversed the Endoplasmic Reticulum (ER) and acquired glycan modifications in the golgi (labeled “post-ER SMO”) is distinguished by its slower migration on the gel. An analysis of additional clones of double mutant *Mgrn1*^-/-^;*Rnf157*^-/-^ cell lines is shown in Fig. S3C. **(B)** Unrooted maximum-likelihood tree topology showing the evolutionary relationship between MGRN1 and RNF157. Branches of the tree are colored according to the major eukaryotic lineages, with the vertebrate-specific RNF157 lineage highlighted in purple. The circle on the node separating MGRN1 and RNF157 denotes 100% confidence support (1000 replicates). The scale bar indicates phylogenetic distance as the number of amino acid substitutions per site. The full Newick tree file is provided in Supplemental File 1. **(C)** Violin plots (left) and corresponding representative confocal fluorescence microscopy images (right) of SMO (red) at primary cilia (green, marked by ARL13B) in wild-type, *Mgrn1*^-/-^, *Mgrn1*^-/-^;*Rnf157*^-/-^, and *Megf8*^-/-^ NIH/3T3 cells (n∼70 cilia/condition). The DNA in the nucleus is stained blue with DAPI. Wild-type cells were analyzed after treatment with either no SHH or saturating SHH (25 nM). Arrowheads identify individual cilia captured in the zoomed images above each panel. Statistical significance was determined by the Kruskal-Wallis test; \*\**p*-value ≤ 0.01 and \*\*\*\**p*-value ≤ 0.0001. Scale bars, 10 µm in merged panels and 2 µm in zoomed displays. An analysis of additional clones of *Mgrn1*^-/-^;*Rnf157*^-/-^ cell lines is shown in Fig. S3D. **(D)** Necropsy (top row) and episcopic confocal microscopy (EFIC, bottom row) images of embryonic hearts from *Mgrn1^m/m^*, *Rnf157^m/m^*, *Megf8^m/m^*, and *Mgrn1^m/m^*;*Rnf157^m/m^* embryos. The type of intracardiac defect (if detected) is noted above the EFIC image. Scale bars, 200 µm. **(E)** Forelimbs of *Megf8^m/m^* and *Mgrn1^m/m^*;*Rnf157^m/m^* embryos show additional preaxial digits (preaxial digit duplication, PDD). Asterisks (*) mark the duplicated preaxial digits. Scale bars, 200 µm. **(F)** Table summarizes the frequency of congenital heart defects (CHDs), heterotaxy, and preaxial digit duplication (PDD) in *Mgrn1^m/m^* (n=15), *Rnf157^m/m^* (n=6), *Megf8^m/m^* (n=12), and *Mgrn1^m/m^*;*Rnf157^m/m^* (n=3) embryos. A detailed list of phenotypes observed in each embryo can be found in Table S1.

These analyses raised the possibility that RNF157 may partially compensate for the loss of MGRN1. Depletion of both RNF157 and MGRN1 in NIH/3T3 cells and NPCs using CRISPR methods (**Fig. S3B**) enhanced Hh signaling activity (**Figs. 1A, S3C-E**). *Mgrn1^-/-^*;*Rnf157^-/-^* NIH/3T3 cells constitutively expressed GLI1, even in the absence of SHH (**Fig. 1A**). In addition, the abundance of SMO carrying mature glycan modifications (acquired in the golgi after trafficking from the endoplasmic reticulum) and the abundance of SMO in primary cilia was much higher in *Mgrn1^-/-^*;*Rnf157^-/-^* compared to *Mgrn1^-/-^* cells (**Figs. 1A and 1C**). In all assays, Hh signaling in *Mgrn1^-/-^*;*Rnf157^-/-^* cells was enhanced compared to *Mgrn1^-/-^* cells (and equivalent to *Megf8^-/-^* cells) (**Figs. 1A and 1C**).

To assess the relationship between RNF157 and MGRN1 *in vivo*, we generated *Rnf157*^-/-^ mice (hereafter referred to as Rnf157*^m/m^* mice) using CRISPR methods (**Fig. S3B**). Consistent with data collected by the International Mouse Phenotyping Consortium (IPMC) using a different knockout strategy (Dickinson et al., 2016), the *Rnf157^m/m^* mice were viable, fertile, and without obvious developmental defects (**Figs. 1D and 1F**). *Rnf157^m/+^* mice were mated to *Mgrn1^m/m^* mice to generate compound heterozygotes, which were intercrossed to obtain *Mgrn1^m/m^*;*Rnf157^m/m^* double null embryos. The penetrance of birth defects in *Mgrn1^m/m^*;*Rnf157^m/m^* embryos was comparable to *Megf8^m/m^* embryos (and much higher than single null *Mgrn1^m/m^* or *Rnf157^m/m^* embryos) (**Figs. 1D-F**). Based on consequences of the simultaneous disruption of both *Rnf157* and *Mgrn1* in both cultured cells and mice, we conclude that RNF157 can partially compensate for the function of MGRN1 in *Mgrn1^-/-^* NIH/3T3 cells, NPCs, and embryos (**Figs. 1A, 1C-1F, and S3C-E**). This compensation is asymmetric, as the loss of RNF157 alone had few developmental consequences (**Figs. 1D and 1F**), presumably because MGRN1 can fully cover RNF157 functions. In conclusion, *Rnf157* is a modifier gene: mutations in *Rnf157* are insufficient to cause a phenotype alone, but they increase the penetrance of phenotypes caused by mutations in a different gene (*Mgrn1*).

### MEGF8 binds to MGRN1

Mouse embryos and cells that lack MEGF8 are indistinguishable from those that lack both MGRN1 and RNF157 (**Figs. 1A and 1C-F**), leading us to speculate that MEGF8, MGRN1, and RNF157 may work together to regulate SMO trafficking. We were encouraged to look for a physical interaction by the BIOGRID and BioPlex databases (Huttlin et al., 2017; Oughtred et al., 2019), which predict that MEGF8 is an interaction partner of both MGRN1 and RNF157 (**Fig. 2A**). To validate this prediction, we transiently expressed MEGF8 in HEK293T cells and observed that it could be co-immunoprecipitated (co-IP) with either endogenous or over-expressed MGRN1 (**Figs. 2A and 2C**). Deleting the ∼170 amino acid (a.a.) long cytoplasmic tail (hereafter called the “Ctail”) of MEGF8 (MEGF8^ΔCtail^), but not its large ∼2500 a.a. extracellular domain (MEGF8^ΔN^), abolished the interaction with MGRN1 (**Figs. 2A-C**). The MEGF8 Ctail contains a peptide motif (with the sequence “MASRPFA”) that is highly conserved across a family of single-pass transmembrane proteins found in Filozoa, animal-like eukaryotes including Filasterea, Choanoflagellatea, and Metazoa (**Figs. 2B and S4A**) (Gunn et al., 1999; Haqq et al., 2003; Nagle et al., 1999). The deletion of this motif (MEGF8^ΔMASRPFA^) abrogated the interaction between MEGF8 and MGRN1 (**Fig. 2C**), establishing an E3 ligase recruitment function for this mysterious sequence element.

**Figure 2:**
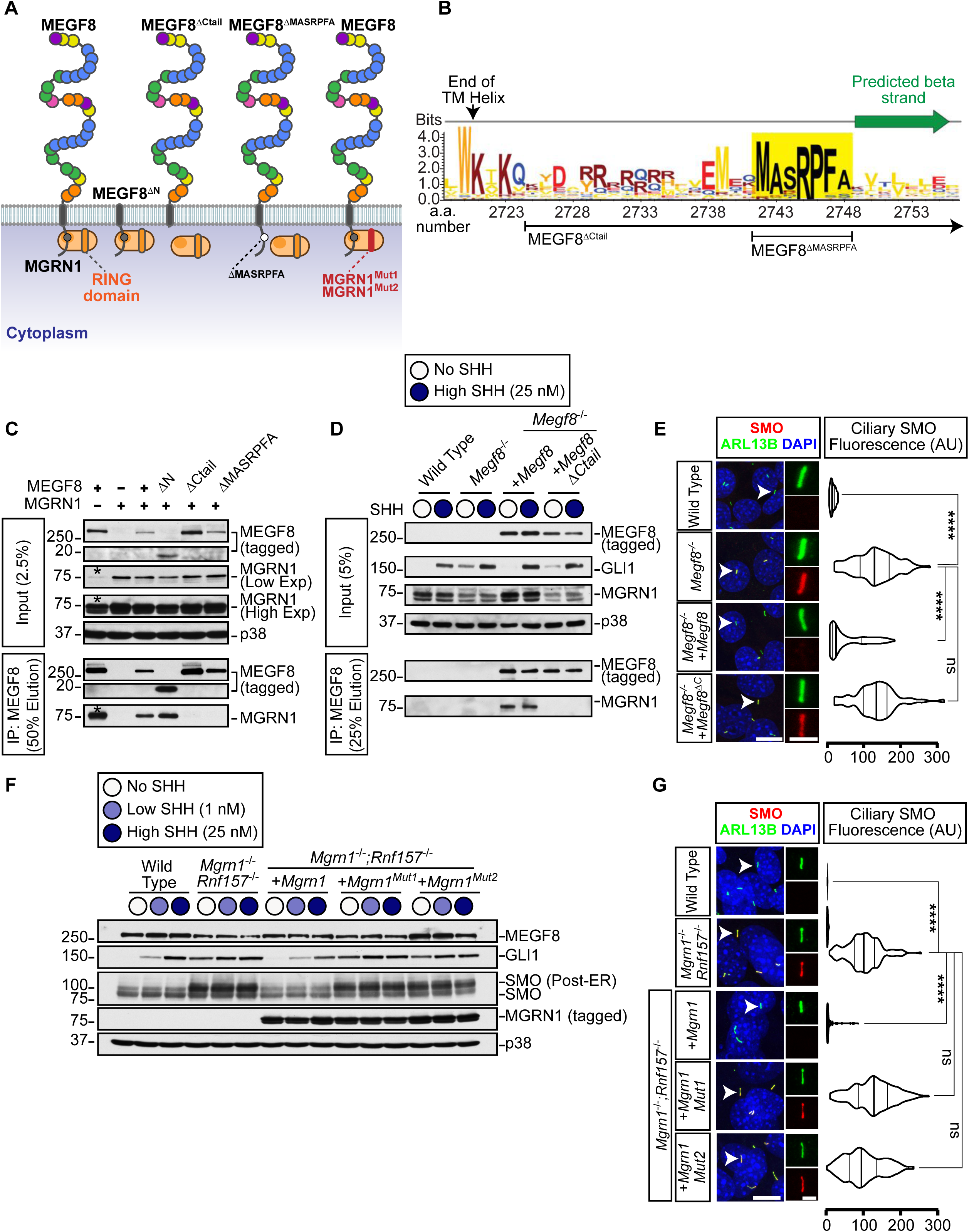
The interaction between MGRN1 and MEGF8 is required to attenuate Hedgehog signaling. **(A)** Depictions of full length MEGF8, truncated MEGF8 (MEGF8^ΔN^, MEGF8^ΔCtail^, MEGF8^ΔMASRPFA^), functional MGRN1, and catalytically inactive MGRN1 (MGRN1^Mut1^ and MGRN1^Mut2^) proteins. The multiple domains in the extracellular region of MEGF8 are shown as circles and colored as in Fig. S1A. **(B)** Sequence logo showing the conservation in sequence entropy bits of the MASRPFA sequence (yellow shading) in the cytoplasmic tail of MEGF8 and related proteins (alignment shown in Fig. S4A). The logo spans the region from the penultimate residue of the TM helix to the extended β-strand-like region immediately distal to the MASRPFA motif (shown by a green arrow). Deletion boundaries for the MEGF8 mutants shown in Fig. 2A are noted below the logo. **(C)** The interaction between MEGF8 or MEGF8 mutants (shown in Fig. 2A, all 1D4 tagged) and MGRN1 (FLAG tagged) was tested by transient co-expression in HEK293T cells, followed by immunoprecipitation (IP) of MEGF8 proteins using the appended 1D4 tag. Immunoblots of input samples show the abundance of indicated proteins in extracts and IP samples show the amount of endogenous and transfected MGRN1 (detected with an anti-MGRN1 antibody) that co-precipitated with MEGF8. Asterisk (*) indicates endogenous MGRN1 from HEK293T cells present in input and IP samples. **(D and E)** GLI1 abundance was measured as a metric of Hh signaling strength by immunoblotting (D) and SMO ciliary abundance by confocal fluorescence microscopy (E) in *Megf8^-/-^* NIH/3T3 cells stably expressing 1D4-tagged MEGF8 or MEGF8^ΔC^ (see Fig. 2A). (D) The interaction between each MEGF8 variant and endogenous MGRN1 was also tested by co-IP. **(F and G)** Total GLI, SMO and MEGF8 abundances were measured by immunoblotting (F) and SMO ciliary abundance by confocal fluorescence microscopy (G) in *Mgrn1^-/-^;Rnf157^-/-^* NIH/3T3 cells stably expressing wild-type MGRN1 or variants carrying inactivating mutations in the RING domain (MGRN1^Mut1^ and MGRN1^Mut2^, see Figs. 2A and S4B). (D, F) Cells used for IP and immunoblotting were treated with the indicated concentrations of SHH; (E, G) cells used for ciliary SMO measurements were left untreated. (E, G) Horizontally positioned violin plots summarize the quantification of SMO fluorescence (red) at ∼50 individual cilia (green, ARL13B) per cell line from representative images of the type shown immediately to the left. (E, G) Arrowheads identify individual cilia captured in the zoomed images to the right of each panel. Statistical significance was determined by the Kruskal-Wallis test; not-significant (ns) > 0.05 and \*\*\*\**p*-value ≤ 0.0001. Scale bars, 10 µm in merged panels and 2 µm in zoomed displays.

To test if the association between MEGF8 and MGRN1 was relevant for the regulation of Hh signaling, we stably expressed wild-type MEGF8 or the interaction-defective MEGF8^ΔCtail^ mutant in *Megf8*^-/-^ NIH/3T3 cells (**Fig. 2D**). Stably expressed MEGF8, but not its truncated MEGF8^ΔCtail^ variant, bound to endogenous MGRN1 (**Fig. 2D**) and suppressed the elevated basal GLI1 and ciliary SMO seen in *Megf8*^-/-^ cells (**Figs. 2D and 2E**). The MEGF8-MGRN1 interaction was unchanged when signaling was activated by the addition of SHH (**Fig. 2D**).

These data establish that MGRN1 in the cytoplasm stably associates with the Ctail of MEGF8 and this interaction is required to suppress ciliary SMO levels and attenuate Hh signaling.

### The ubiquitin ligase activity of MGRN1 is required to attenuate Hh signaling

MGRN1 regulates processes ranging from skin pigmentation to spongiform neurodegeneration by directly ubiquitinating multiple substrates (Chakrabarti and Hegde, 2009; Gunn et al., 2013a; Jiao et al., 2009). We constructed two variants of MGRN1 (MGRN1^Mut1^ and MGRN1^Mut2^) carrying mutations in highly conserved residues of the RING domain (**Fig. S4B**). These mutations are known to abolish binding between RING domains and their cognate E2 partners, thereby preventing ubiquitin transfer to substrates (Garcia-Barcena et al., 2020; Gunn et al., 2013a). We stably expressed wild-type MGRN1, MGRN1^Mut1^, or MGRN1^Mut2^ in *Mgrn1^-/-^*;*Rnf157^-/-^* NIH/3T3 cells and measured the abundance of GLI1, post-ER SMO, and ciliary SMO (**Figs. 2F and 2G**). In all three assays, wild-type MGRN1 was able to fully attenuate Hh signaling and SMO levels, but the MGRN1^Mut1^ and MGRN1^Mut2^ variants were inactive. Importantly, MGRN1^Mut1^ and MGRN1^Mut2^ were expressed at equivalent levels as MGRN1 (**Fig. 2F**) and maintained their stable interaction with MEGF8 (**Fig. S4C**), demonstrating their integrity. These results support the conclusion that both the stable interaction of MGRN1 with MEGF8 and its E3 ligase function are required to attenuate Hh signaling.

### The MEGF8-MGRN1 complex ubiquitinates SMO

At this point our data suggested that MGRN1 functions as a membrane-tethered ubiquitin ligase complex that attenuates Hh signaling by reducing SMO abundance at the cell surface and primary cilium. This mechanism is reminiscent of a prominent membrane-localized ubiquitination system that attenuates WNT signaling by decreasing cell-surface levels of Frizzled (FZD) proteins, receptors for WNT ligands that are the closest relatives of SMO in the GPCR superfamily (Bjarnadóttir et al., 2006). Two transmembrane RING-family E3 ubiquitin ligases, ZNRF3 and RNF43, attenuate WNT responsiveness by directly ubiquitinating FZD and promoting its clearance from the cell surface (Hao et al., 2012; Koo et al., 2012).

To examine if a similar ubiquitination system regulates Hh signaling sensitivity, we measured the stability of SMO at the plasma membrane using a non-cell permeable biotinylation reagent that only labels proteins at the cell surface in wild-type, *Megf8*^-/-^, and *Mgrn1^-/-^*;*Rnf157^-/-^* cells (**Fig. 3A**). Both the steady state abundance and the stability of cell-surface SMO were markedly greater in both mutant cell lines compared to wild-type cells (**Figs. 3B and 3C**). The increase in ciliary SMO abundance (**Fig. 1C**) is likely a secondary consequence of elevated SMO at the plasma membrane, because plasma membrane-localized SMO can enter the cilia by a lateral transport pathway (Milenkovic et al., 2009). These results are analogous to how the stability of cell-surface FZD is enhanced when the ligases ZNRF3 or RNF43 are inactivated (Hao et al., 2012; Koo et al., 2012), prompting us to consider whether SMO is a substrate for the MEGF8-MGRN1 ubiquitin ligase complex.

**Figure 3:**
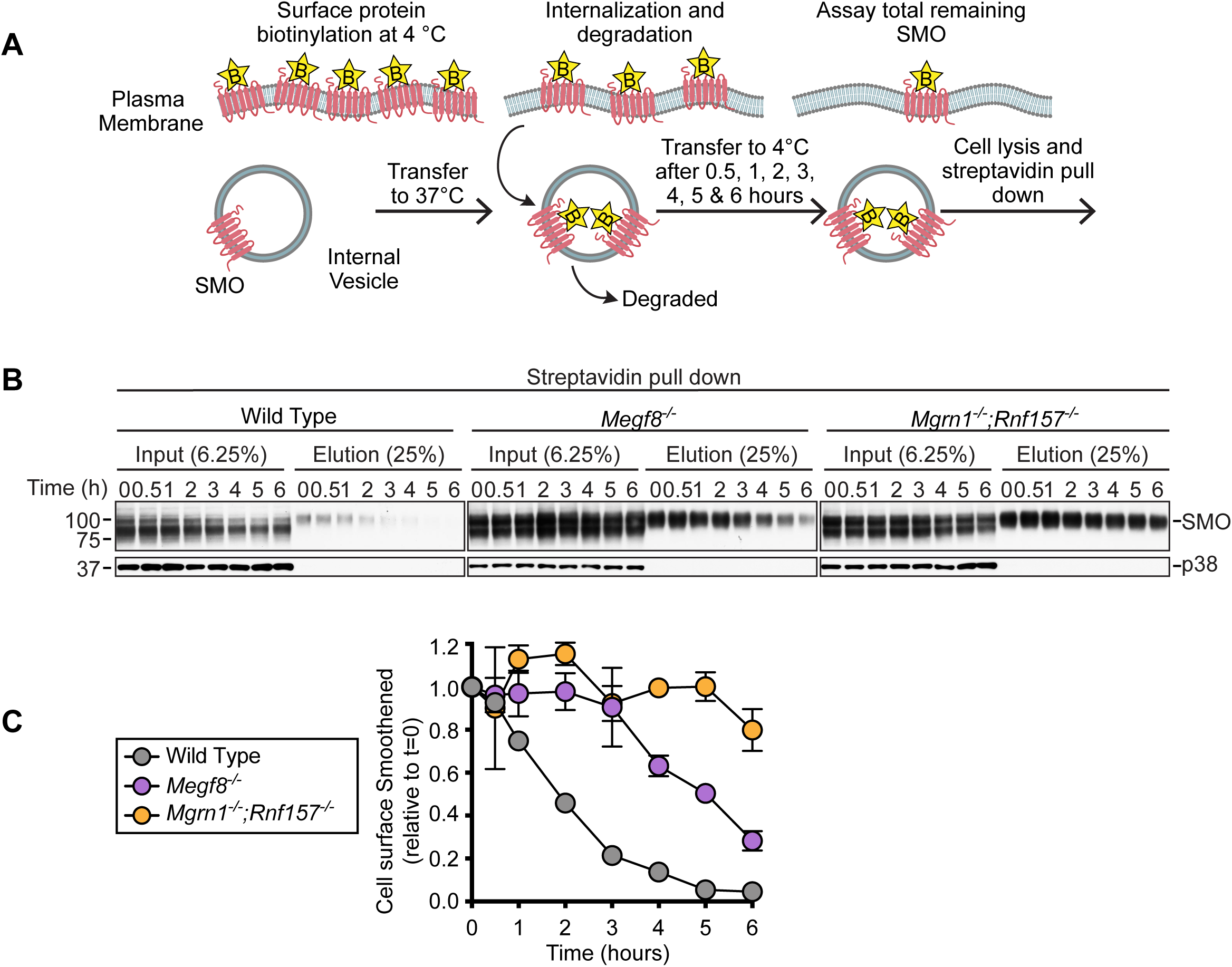
MEGF8 and MGRN1/RNF157 promote the internalization and degradation of SMO at the cell surface. **(A)** To measure the degradation rate of cell-surface SMO in wild-type, *Megf8*^-/-^, and *Mgrn1*^-/-^;*Rnf157*^-/-^ NIH/3T3 cells, SMO was labeled with non-cell-permeable biotin at 4°C. After warming cells to 37°C to allow for SMO internalization and degradation for various periods of time, the amount of biotinylated SMO remaining was isolated on streptavidin beads and measured by immunoblotting **(B)**. While the steady state abundance of SMO (at t=0) was much higher in *Megf8*^-/-^ and *Mgrn1*^-/-^;*Rnf157*^-/-^ cells compared to wild-type cells, the fraction of initial SMO remaining at various times after cell-surface labeling is quantified in **(C)**. Error bars represent the standard error of two independent replicates.

We established an assay to measure SMO ubiquitination by expressing SMO and Hemagglutinin (HA)-tagged ubiquitin (UB) together in HEK293T cells and then measuring the amount of HA-UB conjugated to SMO (**Figs. 4A and 4B**). Lysates were prepared under denaturing conditions to ensure only covalent HA-UB conjugates would survive. SMO was isolated by immunoprecipitation and the attached UB chains detected (as a smear) by immunoblotting with an anti-HA antibody. Co-expression of MGRN1 alone had no effect on SMO ubiquitination, co-expression of MEGF8 alone modestly increased SMO ubiquitination, but the co-expression of both MEGF8 and MGRN1 dramatically increased levels of ubiquitinated SMO and concomitantly reduced SMO abundance (**Fig. 4A**). A ubiquitin mutant lacking all lysine residues (UB^K0^) was poorly conjugated to SMO, suggesting that SMO is attached to poly-UB chains, rather than to a single ubiquitin (**Fig. S5A**). Inactivating mutations in the RING domain of MGRN1 (MGRN1^Mut1^ and MGRN1^Mut2^) failed to promote SMO ubiquitination (**Fig. 4A**), indicating the E3 ligase activity of MGRN1 was required. SMO was a selective substrate for MGRN1 and MEGF8 because their co-expression did not change the abundance of a different ciliary GPCR, SSTR3 (**Fig. S5B**). SMO contains 21 lysine (K) residues exposed to the cytoplasm that could function as acceptors for ubiquitin. Changing all of these lysines to arginines (R) impaired MGRN1-mediated ubiquitination (**Fig. S5C**), but changing specific clusters of lysines in each of the cytoplasmic loops or the tail of SMO did not reduce ubiquitination (**Fig. S5C**). Thus, MGRN1 does not seem to favor a particular lysine residue or set of lysine residues on the cytoplasmic surface of SMO, at least in this over-expression based HEK293T assay.

**Figure 4:**
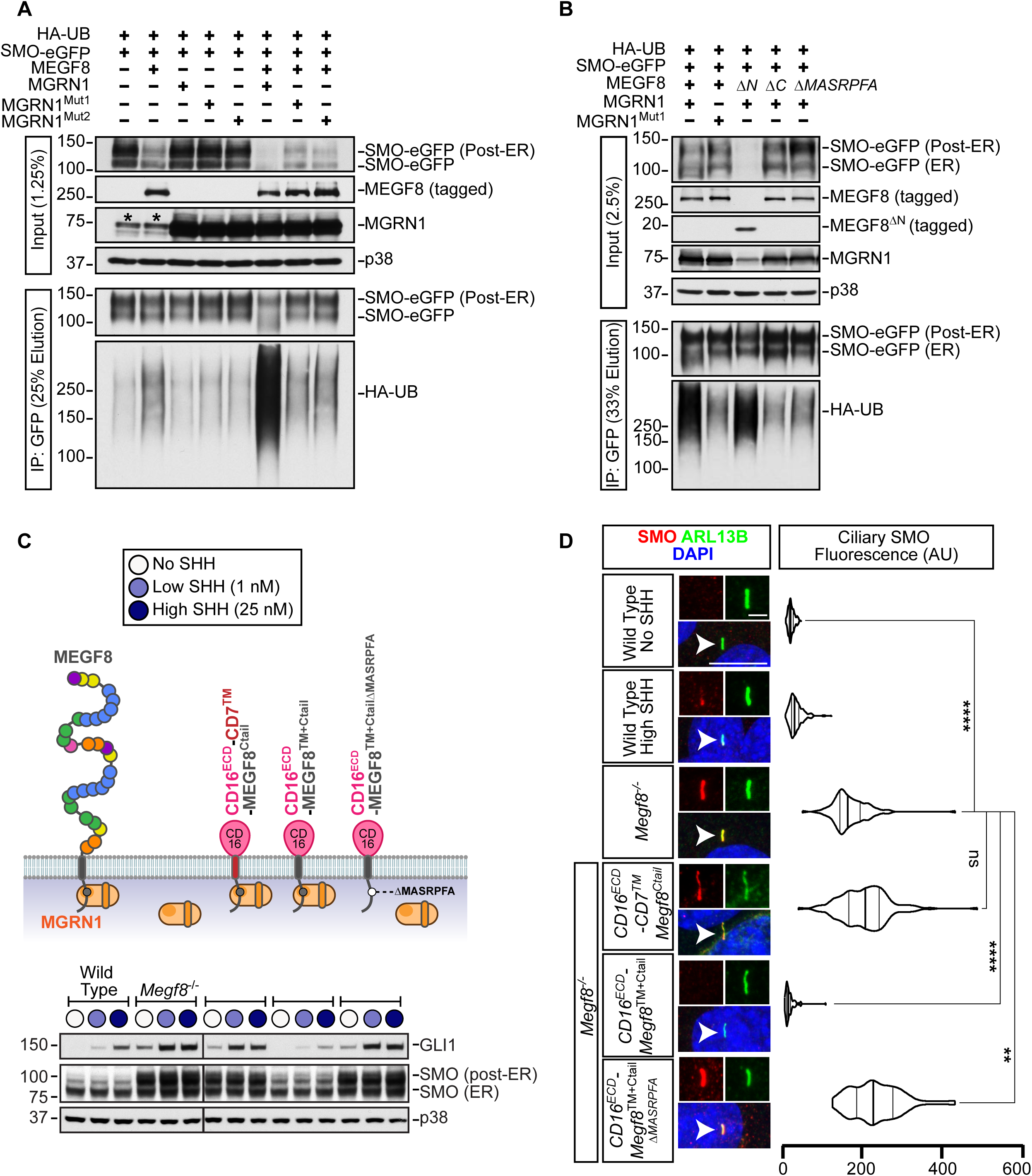
Smoothened is ubiquitinated by the MEGF8-MGRN1 complex. **(A and B)** SMO ubiquitination was assessed after transient co-expression with HA-tagged ubiquitin (HA-UB) and the indicated MEGF8 and MGRN1 variants in HEK293T cells (see Fig. 2A for a summary of protein variants). Cells were lysed under denaturing conditions, SMO was purified by IP, and the amount of HA-UB covalently conjugated to SMO assessed using immunoblotting with an anti-HA antibody. An asterisk (*) indicates endogenous MGRN1 present in HEK293T cells. **(C and D)** Total GLI1 and SMO abundances were measured by immunoblotting (C) and ciliary SMO by fluorescence confocal microscopy (D) in *Megf8^-/-^* cells expressing various CD16/CD7/MEGF8 chimeras (diagrammed in Fig. 4C). Chimeras were composed of the extracellular domain (ECD) of CD16, followed by a transmembrane (TM) helix from either CD7 or MEGF8, and finally by the Ctail of MEGF8, either with or without the MGRN1-interacting MASRPFA motif. The ability of these chimeras to support SMO ubiquitination is shown in Fig. S5D and the abundances of chimeras at the cell surface is shown in Fig. S5E. (D) Horizontally positioned violin plots summarize the quantification of SMO fluorescence (red) at ∼50 individual cilia (green, ARL13B) per cell line from representative images of the type shown immediately to the left. Arrowheads identify individual cilia captured in the zoomed images above each panel. Statistical significance was determined by the Kruskal-Wallis test; not-significant (ns) > 0.05, \*\**p*-value ≤ 0.01, and \*\*\*\**p*-value ≤ 0.0001. Scale bars, 10 µm in merged panels and 2 µm in zoomed displays.

Efficient SMO ubiquitination required both MEGF8 and the E3 ligase function of MGRN1. The small increase in SMO ubiquitination seen in the presence of MEGF8 alone is likely due to presence of endogenous MGRN1 in HEK293T cells (see asterisks in the MGRN1 panel in **Fig. 4A**). To directly test whether the physical interaction between MEGF8 and MGRN1 was required to mediate SMO ubiquitination, we co-expressed MGRN1 with one of three MEGF8 variants (diagrammed in **Fig. 2A**): (1) MEGF8^ΔCtail^, (2) MEGF8^ΔMASRPFA^ (both of which cannot bind to MGRN1, **Fig. 2C**), or (3) MEGF8^ΔN^, which lacks the large extracellular domain of MEGF8 but retains its transmembrane (TM) domain and Ctail. MEGF8^ΔCtail^ and MEGF8^ΔMASRPFA^ failed to support SMO ubiquitination (**Fig. 4B**). In contrast, MEGF8^ΔN^, which can still bind to MGRN1 (**Fig. 2C**), efficiently promoted SMO ubiquitination and degradation (**Fig. 4B**).

Interestingly, MEGF8^ΔN^ was much more active than full-length MEGF8 (despite both proteins being expressed at comparable levels), suggesting that the extracellular domain of MEGF8 may negatively regulate the function of the Ctail or interfere with its interaction with SMO. In addition to recruiting MGRN1 to the plasma membrane, the association between MEGF8 and MGRN1 promoted the intrinsic E3 ligase activity of MGRN1, evident through the ability of MEGF8^ΔN^ to reduce the abundance of co-expressed wild-type MGRN1 (**Fig. 4B**). Most E3 ligases catalyze their own ubiquitination and de-stabilization, a property that reflects their intrinsic catalytic activity.

Unexpectedly, MEGF8^ΔN^, which includes only the TM domain and Ctail of the protein (232 out of the 2778 amino acids in the full-length protein), was sufficient to promote SMO ubiquitination (**Fig. 4B**). To further narrow down the region of MEGF8 required for SMO recognition, we constructed a set of chimeric proteins that fused the MEGF8 Ctail, TM domain, or both to heterologous extracellular and transmembrane domains from CD16 and CD7, respectively (diagrammed in **Fig. S5D**). In the HEK293T assay, both the TM domain and the Ctail of MEGF8 were required to promote SMO ubiquitination; simply tethering the isolated Ctail to the plasma membrane by fusing it to a CD16-CD7 hybrid protein was not sufficient.

Abrogating the interaction with MGRN1 by deleting the “MASRPFA” motif abolished the function of these chimeric proteins, demonstrating that they still require MGRN1 to promote SMO ubiquitination (**Fig. S5D**). If the biochemical function of MEGF8 in Hh signaling is to ubiquitinate SMO, a key prediction is that the CD16^ECD^-MEGF8^TM+Ctail^ chimera, a minimal engineered protein that is sufficient to carry out this function, should be able to reverse the enhanced Hh signaling phenotype in *Megf8*^-/-^ cells. To test this prediction, we stably expressed CD16^ECD^-CD7^TM^-MEGF8^Ctail^, CD16^ECD^-MEGF8^TM+Ctail^, and CD16^ECD^-MEGF8^TM+CtailΔMASRPFA^ (a variant carrying the MASRPFA deletion) in *Megf8*^-/-^ cells (**Fig. 4C**). All three chimeras were expressed and localized properly to the cell surface as measured by flow cytometry using an antibody against the CD16^ECD^ (**Fig. S5E**). However, only the CD16^ECD^-MEGF8^TM+Ctail^ chimera could completely suppress *Gli1* expression and both post-ER and ciliary SMO abundance (**Figs. 4C and 4D**). In these assays, we again observed that the TM domain of MEGF8 was required to regulate Hh signaling, since replacing the TM domain of MEGF8 with the TM domain of CD7 abolished activity (**Figs. 4C and S5D**). We conclude that the TM domain and Ctail of MEGF8 function as a minimal membrane-localized substrate adapter to recruit and activate the E3 ligase activity of MGRN1 towards SMO, catalyzing SMO ubiquitination and clearance from both the plasma and ciliary membrane, and consequently dampening sensitivity to Hh ligands.

### Genetic interactions between *Megf8* and *Mgrn1*

After identifying the MEGF8-MGRN1 interaction and elucidating the ubiquitination based mechanism through which it regulates the sensitivity of target cells to Hh ligands, we sought to investigate the role of this protein complex in embryonic development using the previously published *Megf8^m/+^* and *Mgrn1^m/+^* mouse lines (He et al., 2003; Phillips, 1963; Zhang et al., 2009). Notably, both *Megf8^m/m^* and *Mgrn1^m/m^* mutant embryos display CHDs, heterotaxy, and preaxial digit duplication. While these phenotypes are fully penetrant in the *Megf8^m/m^* mutants, they show lower penetrance in the *Mgrn1^m/m^* mutants (likely due to partial redundancy with *Rnf157*) (**Fig. 1F**) (Cota et al., 2006; Zhang et al., 2009). To determine whether the developmental defects exhibited by these two mutants are a product of the same pathway (as predicted by our biochemical studies), we assessed for a genetic interaction by intercrossing the *Megf8^m/+^* and *Mgrn1^m/+^* mice and examining the phenotypes of the resultant double heterozygous *Megf8^m/+^*;*Mgrn1^m/+^* embryos.

As reported previously (Cota et al., 2006; Zhang et al., 2009), the single heterozygous *Megf8^m/+^* and *Mgrn1^m/+^* embryos were normal without any developmental defects, consistent with the adult viability of *Megf8^m/+^* and *Mgrn1^m/+^* mice (**Figs. 5 and 6**). In contrast, the *Megf8^m/+^*;*Mgrn1^m/+^* double heterozygous embryos showed preaxial digit duplication, heterotaxy and CHDs, phenotypes similar to those seen in homozygous *Megf8^m/m^* and *Mgrn1^m/m^* embryos (**Figs. 5 and 6, Table S4**). Detailed anatomic phenotyping was conducted on e13.5-14.5 *Megf8^m/+^*;*Mgrn1^m/+^* embryos using both necropsies (**Figs. 5B and 5C**) and episcopic confocal microscopy (EFIC) to generate 3D histological reconstructions of the intracardiac anatomy (**Fig. 6B**).

**Figure 5:**
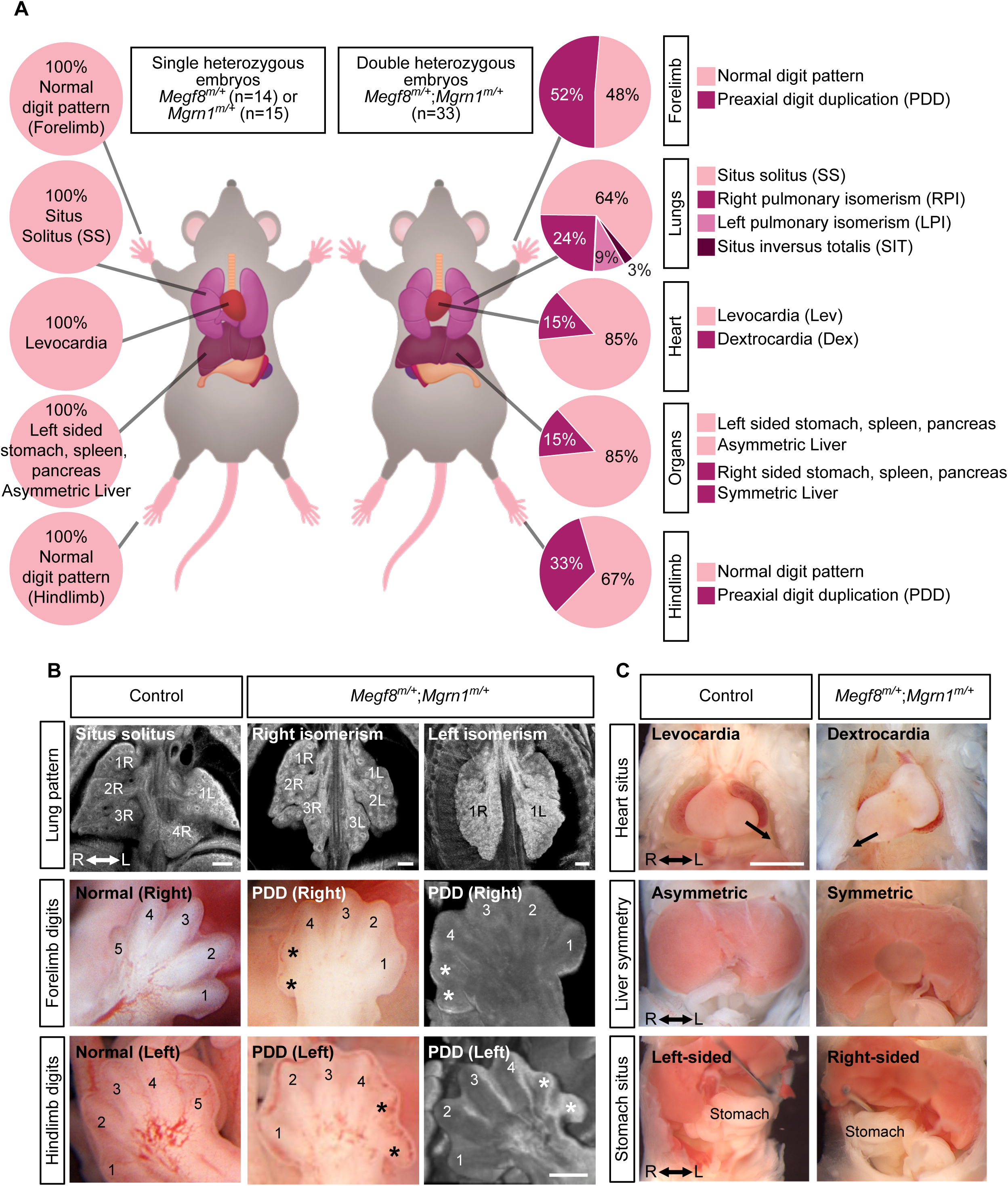
A genetic interaction between *Megf8* and *Mgrn1* causes heart defects and heterotaxy. **(A)** Summary of phenotypes observed in *Megf8*^m/+^ (n=14), *Mgrn1*^m/+^ (n=15), or *Megf8*^m/+^;*Mgrn1*^m/+^ (n=33) mouse embryos (e13.5-14.5). Dex, dextrocardia; Lev, levocardia; LPI, left pulmonary isomerism; PDD, preaxial digit duplication; RPI, right pulmonary isomerism; SIT, situs inversus totalis; SS, situs solitus. A detailed list of phenotypes observed in each embryo can be found in Tables S2, S3, and S4. **(B)** Representative light microscopy and EFIC images of the developing lungs and limbs of single (control) and double heterozygous embryos. The normal right lung has 4 lobes (1R, 2R, 3R and 4R) and the left lung has one lobe (1L). A subset of *Megf8*^m/+^;*Mgrn1*^m/+^ embryos had either right or left pulmonary isomerism. Asterisks (*) in images of the limb mark the duplicated preaxial digits. **(C)** Representative necropsy images showing the position of the heart, symmetry of the liver, and location of the stomach in single (control) and double heterozygous embryos. Arrow (top row) denotes the direction of the cardiac apex, which normally points to the left (levocardia), but occasionally points to the right (dextrocardia) in *Megf8*^m/+^;*Mgrn1*^m/+^ embryos. Liver lobes (middle row) are normally asymmetric, but are occasionally symmetric in *Megf8*^m/+^;*Mgrn1*^m/+^ embryos. The normal location of the stomach (bottom row) is in the left abdomen, but is occasionally present in the right abdomen in *Megf8*^m/+^;*Mgrn1*^m/+^ embryos.

**Figure 6:**
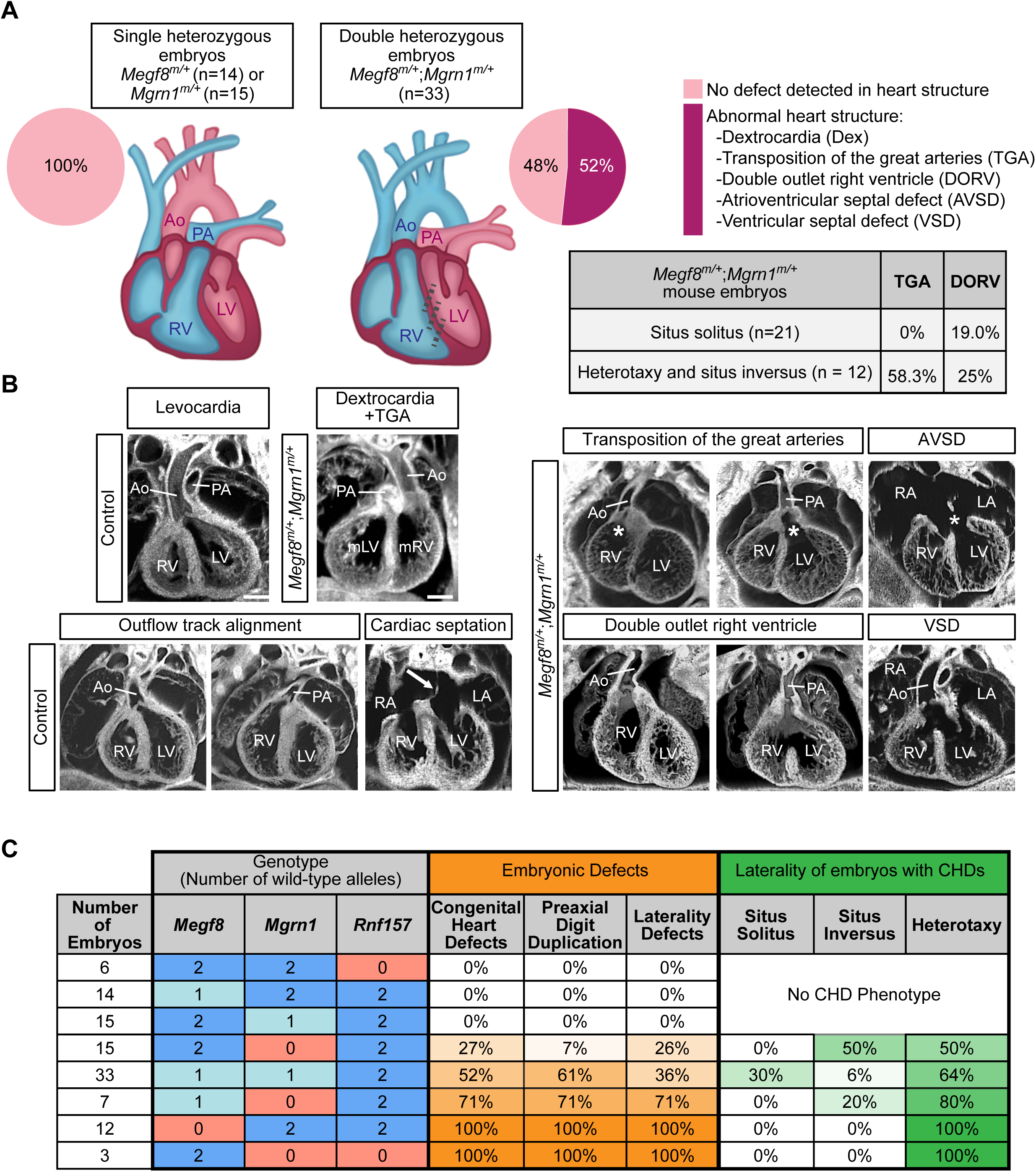
Spectrum of heart defects in mice carrying mutant alleles of *Megf8* and *Mgrn1*. **(A and B)** Summary of congenital heart defects (CHDs) in *Megf8*^m/+^ (n=14), *Mgrn1*^m/+^ (n=15), or *Megf8*^m/+^;*Mgrn1*^m/+^ (n=33) mouse embryos (e13.5-14.5) as determined by EFIC imaging. (A) The table summarizes the frequencies transposition of the great arteries (TGA) and double outlet right ventricle (DORV) in *Megf8^m/+^*;*Mgrn1^m/+^* embryos with or without defects in left-right patterning. (B) Shows representative EFIC images of the many defects observed in double heterozygous embryos, along with normal hearts from control (single heterozygous) embryos. Ao, aorta; AVSD, atrioventricular septal defect; Dex, dextrocardia; LA, left atrium; LV, left ventricle; mLV, morphological left ventricle; mRV, morphological right ventricle; PA, pulmonary artery; RA, right atrium; RV, right ventricle; VSD, ventricular septal defect. A detailed phenotypic analysis of each embryo can be found in Tables S2, S3, and S4. Scale bars, 100 µm. **(C)** Table shows the frequencies of CHDs, preaxial digit duplication, and laterality defects observed in mouse embryos carrying increasing numbers of mutant alleles of *Megf8*, *Mgrn1,* and *Rnf157*. Two wild-type alleles are colored in dark blue, one wild-type allele is colored in light blue, and loss of all functional alleles are colored in light red. Darker shades of orange and green indicate a higher penetrance of the indicated birth defect and laterality phenotype, respectively. A detailed phenotypic analysis of every embryo of each genotype can be found in Tables S1-S5 and a full compilation of the penetrance of various phenotypes is provided in Table S6. For a more detailed analysis of the correlation between laterality and CHD phenotypes observed in Megf8^m/+^;Mgrn1^m/+^ embryos, refer to Table S7.

The limb, heart and left-right patterning defects observed in 100% of *Mefg8^m/m^* embryos (**Table S1**), were incompletely penetrant in *Megf8^m/+^*;*Mgrn1^m/+^* double heterozygous embryos (**Figs. 5 and 6**). Preaxial digit duplication, a hallmark of elevated Hh signaling in the limb bud, was observed in only 61% of *Megf8^m/+^*;*Mgrn1^m/+^* embryos (**Figs. 5A and 6C**). Defects in left-right patterning were seen in only 36% of *Megf8^m/+^*;*Mgrn1^m/+^* embryos (**Figs. 5A and 6C**). Heart defects were seen in ∼52% of *Megf8^m/+^*;*Mgrn1^m/+^* embryos. Only one of the *Megf8^m/+^*;*Mgrn1^m/+^* embryos had *situs inversus totalis,* an inversion of all visceral organs (**Fig. 5A**).

In addition to reduced penetrance, the CHDs seen in *Megf8^m/+^*;*Mgrn1^m/+^* double heterozygous embryos were also milder compared to *Megf8*^m/m^ embryos. All *Megf8*^m/m^ embryos suffered from transposition of the great arteries (TGA), a severe outflow tract (OFT) malalignment defect in which the aorta emerges from the right ventricle and the pulmonary artery from the left ventricle (**Figs. 6A and 6B, Table S1**). Amongst the 52% of *Megf8^m/+^*;*Mgrn1^m/+^* embryos with CHDs, only 41% of these embryos displayed TGA and 47% displayed a milder OFT defect called double outlet right ventricle (DORV) with or without atrioventricular septal defect (AVSD) (**Figs. 6A and 6B, Table S4**).

Given the known co-occurrence of heterotaxy with severe CHDs in clinical data from human birth registries (Lin et al., 2014; Pradat et al., 2003), we examined the correlation between these two types of birth defects in our mutant mouse embryos. All *Megf8*^m/m^ embryos had both heterotaxy and TGA (**Figs. 1F and 6C**, **Table S1**). In *Megf8^m/+^*;*Mgrn1^m/+^* embryos, heterotaxy was associated 100% of the time with CHDs and, conversely, CHDs were associated 64% of the time with heterotaxy (**Fig. 6C**, **Table S4**). Interestingly, the presence of heterotaxy was also correlated with more severe CHDs: ∼60% of these embryos also had TGA (**Fig. 6A**). In contrast, *Megf8^m/+^*;*Mgrn1^m/+^* embryos with normal left-right patterning (*situs solitus*) did not have TGA and instead had the milder DORV in ∼20% of cases (**Fig. 6A**).

These correlations are remarkably similar to data from human birth registries, which report that ∼85% of heterotaxy cases are associated with CHDs that include DORV, TGA and AVSD (Lin et al., 2014; Pradat et al., 2003). The tight association between CHD and heterotaxy is also supported by the observation that all seven embryos with only preaxial digit duplication (but no CHD) had normal *situs solitus* (**Table S4**). Thus, the double heterozygous *Megf8^m/+^*;*Mgrn1^m/+^* embryos recapitulate the known association between severe CHD and heterotaxy seen in human clinical data. The wider spectrum of CHDs seen in these embryos, including DORV, compared to homozygous *Megf8^m/m^* embryos resembles the more diverse range of CHDs seen in human patients with heterotaxy (**Fig. 6A**) (Lin et al., 2014; Pradat et al., 2003).

### Gene dosage effects involving *Mgrn1*, *Megf8* and *Rnf157*

Our comparison of double heterozygous *Megf8^m/+^*;*Mgrn1^m/+^* embryos to homozygous *Megf8^m/m^* embryos suggested that both the penetrance and expressivity of birth defect phenotypes may be determined by precise magnitude of ubiquitin ligase activity, which in turn determines the abundance of SMO and the strength of Hh signaling. This hypothesis predicts that the dosage of *Megf8*, *Mgrn1* and *Rnf157* should influence the penetrance of birth defect phenotypes.

We analyzed embryos carrying varying numbers of loss-of-function *Megf8^m^*, *Mgrn1^m^*, and *Rnf157*^m^ alleles (**Fig. 6C**). *Megf8^m/m^* and *Mgrn1^m/m^;Rnf157^m/m^* embryos have a 100% penetrance of CHDs, heterotaxy, and preaxial digit duplication, presumably because the functions of both the transmembrane adaptor (MEGF8) and the cytoplasmic E3 ligases (MGRN1 or RNF157) are essential for SMO ubiquitination. Loss of one allele of *Megf8* (*Megf8^m/+^* embryos), one allele of *Mgrn1* (*Mgrn1^m/+^* embryos) or both alleles of *Rnf157* (*Rnf157^m/m^* embryos) did not lead to birth defects, likely because the abundance of the MEGF8-MGRN1/RNF157 complex remains above the threshold required for normal development.

However, between these two extremes, decreasing the cumulative gene dosage (by increasing the number of mutant alleles) of *Mgrn1* and *Megf8* led to a progressive increase in the penetrance of CHDs, heterotaxy and preaxial digit duplication (**Fig. 6C**). In addition, the incidence of TGA (**Table S6**), the most severe CHD, and the co-occurance of heterotaxy (**Fig. 6C**) increased with decreasing gene dosage. These striking gene dosage effects support the model that a progressive decrease in ubiquitin ligase function leads to a progressive increase in the penetrance and expressivity of birth defects, perhaps by driving a graded increase in Hh signaling strength.

The exquisite sensitivity of heart development to mutations in *Megf8*, *Mgrn1* and *Rnf157* seen in mouse embryos prompted us to look for potentially damaging variants in these genes in patients with CHDs. Using whole exome sequencing data from a cohort of 652 CHD patients, we searched for missense variants in all three genes with a Combined Annotation Dependent Depletion (CADD) score >10. We additionally used a stringent mean allele frequency (MAF) filter of < 0.5% for *MEGF8* and *MGRN1*, but a more relaxed MAF filter (< 5%) for *RNF157*, since the *Rnf157^m/m^* mouse has no phenotype. Using these criteria, we identified one patient (7501) with two mutations each in *MEGF8* and MGRN1 and one mutation in RNF157 (**Figs. S6A and S6B**). Genotyping the parents of patient 7501 revealed that the two mutations in MEGF8 and MGRN1 were both present in the same allele, with the former transmitted from the mother and the latter from the father (along with the RNF157 variant). Patient 7501 clinically presented with OFT anomalies: pulmonary atresia (**Fig. S6C**), a severely hypoplastic right ventricle with an intact interventricular septum (**Fig. 7B**), an atrial septal defect (**Fig. 7B**) and patent foramen ovale (**Fig. 7B**). Primary fibroblasts from patient 7501 displayed increased abundance of ciliary SMO (**Fig. 7C**) and elevated *Gli1* expression (**Fig. 7D**), both at baseline and in response to SHH, when compared to fibroblasts generated from a subject without CHD. Collectively, our mouse and human data support a model where disruption of the MEGF8-MGRN1/RNF157 ubiquitin ligase complex can lead to elevated SMO, increased Hh signaling strength and, consequently, to the emergence of CHDs.

**Figure 7:**
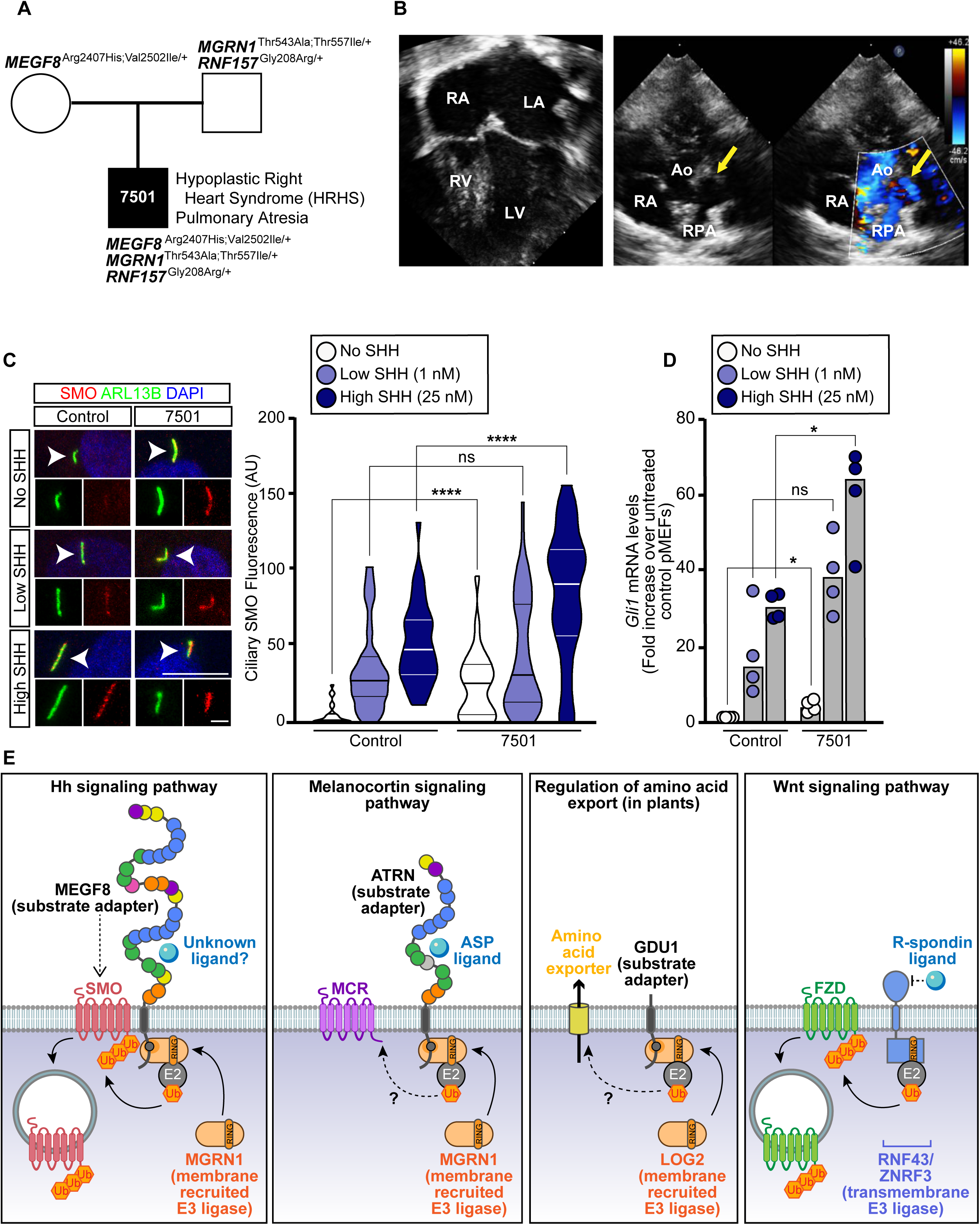
Damaging variants in *MEGF8*, *MGRN1*, and *RNF157* are associated with congenital heart defects in humans. **(A)** Trio pedigree analysis showing the inheritance of *MEGF8*, *MGRN1*, and *RNF157* variants and the presence of severe CHDs. Unaffected parents have variants in either *MGRN1 and RNF157* (father) or *MEGF8* (mother), while the affected progeny (referred to as patient 7501) inherited variants in both genes. The position of these variants in *MEGF8*, *MGRN1*, and *RNF157*, their evolutionary conservation, allele frequency and predicted damaging effect on protein function are shown in Figs. S6A and S6B. Whole exome sequencing results from patient 7501 can be found in Table S8. **(B)** (Left) Four-chamber view of echocardiogram from patient 7501 demonstrating a hypoplastic right ventricle (RV). (Right) Short axis view of the echocardiogram from patient 7501 demonstrated membranous pulmonary atresia (yellow arrow). RA, right atrium; LA, left atrium; LV, left ventricle; RPA, Right Pulmonary Artery; Ao, Aorta. **(C)** Confocal fluorescence microscopy was used to measure the abundance of ciliary SMO (red) at cilia (green, ARL13b) after treatment with no, low (1 nM), or high (25 nM) concentrations of SHH in primary fibroblasts from patient 7501 or a control individual with no known CHDs or variants of interest. Arrowheads identify individual cilia captured in the zoomed images below each panel. Scale bars are 10 µm in merged panels and 2 µm in zoomed displays. The violin plot on the right summarizes the quantification of SMO at ∼20-50 cilia for each condition. Statistical significance was determined by the Mann-Whitney test; not-significant (ns) > 0.05 and \*\*\*\**p*-value ≤ 0.0001. **(D)** Hh signaling strength was assessed using qRT-PCR to measure the abundance of *Gli1* mRNA in primary fibroblasts from patient 7501 and an unaffected control. Bars denote the median *Gli1* mRNA values derived from the four individual measurements shown. Statistical significance was determined by the Mann-Whitney test; not-significant (ns) > 0.05 and \**p*-value ≤ 0.05. **(E)** Regulation of signaling and transport by receptor-like E3 ubiquitin ligases. A model for the mechanism of SMO regulation by the MEGF8-MGRN1 complex (far left) highlights its conceptual similarity to the regulation of melanocortin receptors (MCRs) by the ATRN-MGRN1 complex (middle left), amino acid export by the GDU1-LOG2 complex in plants (middle right), and Frizzled (FZD) receptors for WNT ligands by the ZNRF3/RNF43 family of transmembrane E3 ligases (far right). MEGF8 functions as a transmembrane substrate adaptor, recruiting MGRN1 (and presumably an unknown E2 enzyme) through its cytoplasmic tail to promote the ubiquitination of SMO. SMO ubiquitination leads to its internalization and degradation, thus attenuating responses to Hh ligands. Note that the RING superfamily E3 ligase domains in ZNRF3 and RNF43 are directly fused to a transmembrane and extracellular domain that binds to R-Spondin ligands. We speculate that MEGF8 activity may be regulated by a luminal or extracellular signal (labeled unknown ligand).

## Discussion

Using a combination of mechanistic studies, mouse genetics, and deep anatomical phenotyping, we uncovered a unique membrane-tethered ubiquitination pathway that regulates developmental patterning in a variety of tissues by controlling the trafficking of signaling receptors. MEGF8 functions as a transmembrane substrate adaptor that recruits a cytoplasmic E3 ligase (MGRN1) to catalyze the ubiquitination of SMO, leading to its endocytosis and degradation (**Fig. 7E**). This ubiquitination reaction reduces the abundance of SMO at the cell surface and primary cilium and, consequently, dampens Hh signaling in target cells.

### Receptor-like ubiquitin ligases attenuate signaling strength

The architecture of the MEGF8-MGRN1 complex is notable for the presence of a membrane-spanning component with an extracellular or luminal domain (**Fig. 7E**). This feature suggests a receptor-like function, conceptually analogous to receptor kinases, to transmit extracellular or luminal signals across the membrane to alter the ubiquitination of substrates in the cytoplasm. Interestingly, FZD receptors for WNT ligands are regulated by transmembrane E3 ligases (RNF43 and ZNRF3) in which the RING-containing domain is directly fused to the membrane-spanning component (rather than being non-covalently associated as in the MEGF8-MGRN1 system) (**Fig. 7E**). While a ligand for MEGF8 remains unknown, ZNRF3 and RNF43 are regulated by R-Spondin ligands, critical regulators of progenitor cells during development and stem cells in adult tissues (Hao et al., 2012; Koo et al., 2012). The ubiquitination of receptors by membrane-tethered E3 ligases represents an attractive post-transcriptional mechanism to control the sensitivity of tissues to signaling ligands during development or tissue renewal.

Evolutionary sequence analysis supports a widespread (but under-appreciated) role for transmembrane E3 ligase complexes in ubiquitin signaling. In animals and their immediate sister lineages, MGRN1 and RNF157 likely function as common components of multiple membrane-tethered E3 ligase complexes featuring members of the MEGF8 family of cell-surface proteins, all of which contain an equivalent of the cytoplasmic MASRPFA motif (**Fig. S4A**) (Gunn et al., 1999; Haqq et al., 2003; Nagle et al., 1999). For example, MGRN1 and a different member of this family, Attractin (ATRN), have been implicated in regulation of melanocortin receptor levels by ubiquitination (**Fig. 7E**) (Cooray et al., 2011; Walker, 2010). However, members of the MGRN1 family of RING finger proteins are more widely distributed across eukaryotes (**Fig. 1B**) compared to MEGF8/ATRN-like proteins (which are restricted to animal-like eukaryotes) (Pusapati et al., 2018b). Thus, MGRN1-family proteins may associate with other adaptors beyond the MEGF8/ATRN family to form comparable membrane-localized E3 ligase complexes in different eukaryotic lineages. Indeed, a plant ubiquitin ligase, LOG2, which belongs to the MGRN1 family, associates with and ubiquitinates a single TM protein Glutamine dumper-1 (GDU1) which in turn regulates amino acid transport (**Fig. 7E**) (Guerra et al., 2013). Strikingly, human MGRN1 can functionally replace LOG2 in plants (Guerra et al., 2013). Like the MEGF8/ATRN family in animals, the extensive GDU1 family, conserved across all land plants, has a cytoplasmic tail which features a conserved motif. This motif has a central “MAs” signature (where s is a small amino acid typically G or S) reminiscent of the MASRPFA sequence in the MEGF8/ATRN family (**Fig. 2B**). Accordingly, we propose that the MGRN1 family of RING E3 ligases can associate more generally across eukaryotes with single-pass TM proteins with conserved cytoplasmic motifs, each of which function as a substrate adaptor to target the ubiquitination of specific receptors or transporters (**Fig. 7E**).

### Role of Hh signaling in left-right patterning and heart development

Several lines of evidence point to the Hh signaling pathway as being important for the phenotypes seen in mice carrying mutant alleles of *Megf8* and *Mgrn1*. Both genes were identified in our unbiased genetic screen as attenuators of Hh signaling (Pusapati et al., 2018a) and the key Hh transducer SMO is a direct substrate of the MEGF8-MGRN1 complex. Hh signaling plays a key role in each of the developmental events where we see prominent birth defect phenotypes: left-right patterning, cardiac morphogenesis, and limb development.

Hh signaling has previously been implicated in sustaining left-right patterning signals, a very early event in development that directs the correct asymmetric development of the heart and other visceral organs (Levin et al., 1995; Tsiairis and McMahon, 2009; Zhang et al., 2001). The genetic deletion of SMO, which reduces Hh signaling strength, disrupts left-right patterning and causes a midline heart tube that fails to loop to the right and an embryo that fails to turn (Zhang et al., 2001). Distinct from prior work showing that the loss of Hh signaling impairs proper left-right patterning in the mouse, we find that mutations (in *Megf8* and *Mgrn1*) that *increase* Hh signaling strength also result in heterotaxy and CHDs. These results support the view that left-right patterning (and cardiac and limb development) depend on a just-right “goldilocks” level of Hh signal amplitude or duration.

Hh signaling also influences multiple aspects of heart development: development of the secondary heart field and proper septation of the atria and outflow tract (Dyer and Kirby, 2009; Hoffmann et al., 2009; Washington Smoak et al., 2005). We expect that CHDs seen in our mutant mice are caused by both heterotaxy and by later defects in Hh-mediated patterning of the cardiac septa and outflow tract. The common link of both processes to precisely-tuned level of Hh signaling may explain the tight association between heterotaxy and CHDs that has been long-noted in clinical studies and is recapitulated in our mutant mouse embryos (Pradat et al., 2003).

We acknowledge that MEGF8 and MGRN1/RNF157 may regulate signaling receptors other than SMO and some of the birth defect phenotypes we observe may be related to disruption of other signaling pathways. Future priorities include the identification of other substrates targeted for ubiquitination by the MEGF8-MGRN1 complex and the assessment of where and when these proteins function during embryonic development.

### Oligogenic interactions and gene dosage effects underlie birth defects

Our biochemical studies provide an explanation for the strong genetic interactions observed between the genes that encode subunits of the MEGF8-MGRN1/RNF157 complex. While single heterozygous *Megf^m^*^/+^ and *Mgrn1*^m/+^ embryos are normal, double heterozygous *Megf^m^*^/+^;*Mgrn1*^m/+^ embryos display CHDs with heterotaxy. This phenomenon has been called “synthetic haploinsufficiency” and can result in an oligogenic pattern of inheritance, where mutations in one gene affect the phenotypic outcome of mutations in a different gene (Kousi and Katsanis, 2015; Veitia et al., 2013). Synthetic haploinsufficiency is most commonly seen between genes that encode subunits of a protein complex, like MEGF8 and MGRN1 (Veitia, 2010). Pioneering studies of Bardet-Biedl Syndrome (BBS) and other inherited retinopathies have demonstrated the importance of oligogenic interactions for understanding the genetic etiology of human diseases (Badano et al., 2006; Katsanis et al., 2000).

Beyond binary genetic interactions, the penetrance and expressivity of birth defect phenotypes progressively increases as an inverse function of the gene dosage of *Megf8*, *Mgrn1* and *Rnf157.* We propose that this quantitative effect of mutations in this pathway is explained by the dependence of proper left-right patterning and cardiac development on the precise amplitude of Hh signaling and by the central role of the MEGF8-MGRN1 pathway in setting this amplitude in target cells. The inheritance of increasing numbers of *Megf8*, *Mgrn1* and *Rnf157* mutant alleles will lead to a progressive decrease in the abundance (and hence activity) of the MEGF8-MGRN1/RNF157 complex. Decreasing E3 ligase activity will result in progressive increases in cell surface and ciliary SMO and thus increases in target cell sensitivity to Hh ligands. More generally, our results show that developmental patterning events can be tightly regulated by mechanisms in target cells that function to precisely tune sensitivity to extracellular morphogens.

We finish by noting that our genetic analyses highlight how interactions between a small number of genes can produce a complex inheritance pattern (common to many human diseases) that features sporadic occurrence, incomplete penetrance, and variable expressivity. Homozygous mutations in *Megf8* result in a uniform phenotypic spectrum, with 100% of embryos displaying TGA, heterotaxy, and preaxial digit duplication. However, the co-inheritance of one mutant allele of *Megf8* with one mutant allele of *Mgrn1* (even in the homogenous genetic background of inbred mice) results in both incomplete penetrance and variable expressivity of phenotypes, manifested by a wider range of CHDs like TGA, DORV, and septal defects. Indeed, whole-exome sequencing studies of human CHD cohorts increasingly support a prominent role for such oligogenic inheritance mechanisms in the genetic etiology of CHDs (Gifford et al., 2019; Jin et al., 2017; Liu et al., 2017, 2020; Priest et al., 2016).

## Acknowledgements

We thank Derek Silvius and Janet Peters for genotyping and the McLaughlin Research Institute Animal Resource staff for animal care. We thank Connie Chuy for her mouse and heart diagrams. RR was supported by a grant from the National Institutes of Health (GM118082),GP by a postdoctoral fellowship from the American Heart Association (14POST20370057), and JK by a postdoctoral fellowship from the American Heart Association (19POST34380734) and a K99/R00 award from the NIH (GM13251801). CWL, CBY, and SH were supported by grants from the NIH (HL142788 and HL132024) and the DOD (W81XWH-15-1-0649 and W81XWH-16-1-0613).

## Author contributions

Conceptualization, JHK, GVP, RR, CWL, TMG; Methodology, JHK, CBY, TMG; Formal analysis, AK, LA, JHK, CBY; Investigation, JHK, GVP, CBY, TMG, CP, SH, JIL, WD, AMB, TSA; Resources, RR, CWL, TMG, JIL, WD; Writing--Original Draft, JHK, RR, CWL; Writing-Revision and Editing, JHK, RR, CWL, TMG, GVP, CBY; Visualization, JHK, GVP, CBY, AK, LA, CBY; Supervision, RR, CWL, TMG, LA; Project Administration, JHK; Funding Acquisition, RR, CWL, TMG, LA.

## STAR Methods

### LEAD CONTACT AND MATERIALS AVAILABILITY

Further information and requests for resources and reagents should be directed to and will be fulfilled by the Lead Contact, Rajat Rohatgi. All unique/stable reagents generated in this study are available from the Lead Contact with a completed Materials Transfer Agreement.

### EXPERIMENTAL MODEL AND SUBJECT DETAILS

#### NIH/3T3 and HEK293T cell culture

Flp-In-3T3 (a derivative of NIH/3T3 cells and referred to as “NIH/3T3” cells throughout the text) and HEK293T cell lines were purchased from Thermo Fisher Scientific and ATCC, respectively. Information on the gender of the cell lines is not available. NIH/3T3 and HEK293T cells were cultured in Complete Medium, which contained: Dulbecco’s Modified Eagle Medium (DMEM) containing high glucose (Thermo Fisher Scientific, Gibco) and supplemented with 10% fetal bovine serum (FBS) (MilliporeSigma), 2 mM L-Glutamine (Gemini Bioproducts), 1 mM sodium pyruvate (Thermo Fisher Scientific, Gibco), 1x MEM non-essential amino acids solution (Thermo Fisher Scientific, Gibco), and penicillin (40 U/ml) and streptomycin (40 µg/ml) (Gemini Bioproducts). The NIH/3T3 and HEK293T cells were passaged with 0.05% Trypsin/EDTA (Gemini Bioproducts). All cells were housed at 37 °C in a humidified atmosphere containing 5% CO_2_. Cell lines and derivatives were free of mycoplasma contamination as determined by PCR using the Universal Mycoplasma Detection Kit (ATCC).

### Generation of primary mouse embryonic fibroblasts

Primary mouse embryonic fibroblasts (pMEFs) were generated using a modified published protocol (Durkin et al., 2013). Briefly, e12.5-14.5 embryos were harvested and rinsed thoroughly with PBS to remove any excess blood. Using forceps, the head and internal organs (heart and liver) were removed. The embryos were then separated into individual dishes and a sterile razor blade was used to physically mince the tissue in 0.25% Trypsin/EDTA (Thermo Fisher Scientific, Gibco). After pipetting the minced tissue up and down several times to further break up the tissue, the dishes were placed in a 37 °C tissue culture incubator for 10-15 minutes. If there were still large tissue pieces present, the minced tissue was pipetted further and the dish was placed in the incubator for an additional 5-10 minutes. The trypsin was then deactivated using Complete Medium (containing 10% FBS). The cells were then centrifuged, resuspended in fresh Complete Medium, and plated. Each clonal cell line represents pMEFs generated from a single embryo. The gender of the embryos were not determined prior to generating the pMEF cultures. Cells were housed at 37 °C in a humidified atmosphere containing 5% CO_2_.

### Patient recruitment and nasal sampling for patient derived fibroblast cultures

Patients and parents were recruited from the Children’s Hospital of Pittsburgh with informed consent obtained under a human study protocol approved by the University of Pittsburgh Institutional Review Board. Control, CHD patient, and parents recruited had blood drawn for DNA extraction. CHD diagnosis was confirmed with examination of the patient’s medical records. Nasal tissue was obtained from the patient by curettage of the inferior nasal turbinate using a rhino probe. The nasal epithelial tissue was plated in RPMI medium (Thermo Fisher Scientific, Gibco) with 10% FBS (MilliporeSigma) and the fibroblast outgrowths that emerged were expanded and used for Hh signaling assays (below). Both primary fibroblast cell lines (control and patient 7501) were derived from cells collected from female patients.

### Hh signaling assays in NIH/3T3 cells and primary fibroblasts

For Hh signaling assays, NIH/3T3 cells, pMEFs, and primary human fibroblasts were first grown to confluence in Complete Medium (containing 10% FBS) and then ciliated by changing the cell medium to Low Serum Medium (complete medium containing 0.5% FBS) for 24 hours. Cells were then treated with either no SHH, a low concentration of SHH (1 nM), a high concentration of SHH (25 nM), or SAG (100 nM) for at least 4 hours prior to fixation (for NIH/3T3 immunofluorescence assays), 24 hours prior to lysis (for NIH/3T3 Western blot assays), or 48 hours prior to experimentation (for pMEF and primary human fibroblast immunofluorescence and western blot assays).

Hh signaling activity was measured using real-time quantitative reverse transcription PCR (qRT-PCR). RNA was extracted from NIH/3T3 cells and mouse pMEFs using TRIzol reagent (Thermo Fisher Scientific, Invitrogen) as previously described (Rio et al., 2010). Equal amounts of RNA were used as template for cDNA synthesis using the iScript Reverse Transcription Supermix (Bio-Rad Laboratories). qRT-PCR for m*Gli1* and m*Gapdh* was performed on a QuantStudio 5 Real-Time PCR System (Thermo Fisher Scientific) with the following custom designed primers: m*Gli1* (Fwd 5’-CCAAGCCAACTTTATGTCAGGG-3’ and Rev 5’-AGCCCGCTTCTTTGTTAATTTGA-3’) and m*Gapdh* (Fwd 5’-AGTGGCAAAGTGGAGATT-3’ and Rev 5’-GTGGAGTCATACTGGAACA-3’). Similarly, RNA was isolated from primary human fibroblasts using the RNeasy Plus Mini Kit (Qiagen). Equal amounts of RNA were used as template for human cDNA synthesis using the High-Capacity RNA-to-cDNA Kit (Thermo Fisher Scientific, Applied Biosystems). qRT-PCR for h*GLI1* and h*GAPDH* was performed on a 7900HT Real-Time PCR System (Life Technologies) with the following primers: *hGLI1* (Fwd 5’-CAGGGAGGAAAGCAGACTGA-3’ and Rev 5’-ACTGCTGCAGGATGACTGG-3’) and *hGAPDH* (Fwd 5’-GTCTCCTCTGACTTCAACAGCG-3’ and Rev 5’-ACCACCCTGTTGCTGTAGCCAA-3’). For all qRT-PCR experiments, *Gli1* transcript levels were calculated relative to *Gapdh* and reported as a fold change across conditions using the comparative C_T_ method (ΔΔC_T_ method).

### Neural progenitor differentiation assay

Maintenance of HM1 mouse embryonic stem cells (mESCs) harboring the GLI-Venus and OLIG2-mKate dual reporter system and their differentiation into neural progenitor cells (NPCs) was performed as described previously (Pusapati et al., 2018a). The parental HM1 mESC line was derived from a male mouse. mESCs were grown and maintained on feeder cells in mESC Medium, which contains: Dulbecco’s Modified Eagle Medium (DMEM) containing high glucose (Thermo Fisher Scientific, Gibco) and supplemented with 15% FBS (MilliporeSigma), 2 mM L-Glutamine (Gemini Bioproducts), 1 mM sodium pyruvate (Thermo Fisher Scientific, Gibco), 1x MEM non-essential amino acids solution (Thermo Fisher Scientific, Gibco), 1% penicillin/streptomycin (Gemini Bioproducts), 1% EmbryoMax nucleosides (MilliporeSigma), 55 µM 2-mercaptoethanol (Thermo Fisher Scientific, Gibco), and 1000 U/ml ESGRO LIF (MilliporeSigma). mESCs were differentiated into spinal neural progentiors using a previously described protocol (Sagner et al., 2018). mESCs were panned to clear the feeder cells, then plated on 6-well gelatin-coated CellBIND plates (Corning) at a density of 100,000 cells/well. Differentiation was conducted in N2B27 Medium which contains: DMEM/F12 (Thermo Fisher Scientific, Gibco) and Neurobasal medium (Thermo Fisher Scientific, Gibco) (1:1 ratio) supplemented with 1x N-2 supplement (Thermo Fisher Scientific, Gibco), 1x B-27 supplement (Thermo Fisher Scientific, Gibco), 1% penicillin/streptomycin (Gemini Bioproducts), 2 mM L-Glutamine (Gemini Bioproducts), 55 µM 2-mercaptoethanol (Thermo Fisher Scientific, Gibco), and 40 µg/ml bovine serum albumin (MilliporeSigma). On Day 0 (the day the cells are plated) and Day 1, the N2B27 medium was supplemented with 10 ng/ml bFGF (R&D Systems). On Day 2, the N2B27 medium was supplemented with 10 ng/ml bFGF (R&D Systems) and 5 µM CHIR 99021 (Axon Medchem). On Day 3, the N2B27 medium was supplemented with 100 nM Retinoic Acid (RA) (MilliporeSigma) and either no SHH, 5 nM (low SHH), or 25 nM (high SHH). The cells were cultured in RA and SHH for a total of 3 days, where the medium was changed every 24 hours. On Day 6, the cells were washed with PBS and trypsinized with 0.25% Trypsin/EDTA (Thermo Fisher Scientific, Gibco) for flow cytometry analysis. GLI-Venus and OLIG2-mKate fluorescence was measured on a FACScan Analyzer at the Stanford Shared FACS Facility. To detect GLI-Venus, a 488 nm (blue) laser was used with a 525/50 filter and B525 detector. To detect OLIG2-mKate, a 561 nm (yellow) laser was used with a 615/25 filter and Y615 detector.

### Generation of knockout cell lines

Clonal *Mgrn1^-/-^* NIH/3T3 lines were previously generated using a dual single guide (sgRNA) strategy and validated (Pusapati et al., 2018a). Clonal double knockout *Mgrn1^-/-^;Rnf157^-/-^* NIH/3T3 lines were generated using the same dual sgRNA strategy to target *Rnf157* in *Mgrn1^-/-^* NIH/3T3 cells. Briefly, sgRNAs targeting *Rnf157* were designed using the Broad Institute Genetic Perturbation Platform sgRNA Designer Tool (https://portals.broadinstitute.org/gpp/public/analysis-tools/sgrna-design): Exon 6, 5’-CCACAGCGTGCACTACCAGA-3’ and Exon 7, 5’-CAAAAGTGCCCAGAAGCACG-3’. The sgRNAs were then were cloned into pSpCas9(BB)-2A-GFP (Addgene) (Ran et al., 2013) and pSpCas9(BB)-2A-mCherry (Pusapati et al., 2018a) and transfected into NIH/3T3 cells using X-tremeGENE 9 DNA transfection reagent (Roche Molecular Systems). Five days post transfection, GFP and mCherry double positive single cells were sorted into a 96-well plate using a FACSAria II at the Stanford Shared FACS Facility. To detect the GFP, a 488nm (blue) laser was used with a 530/30 filter and B530 detector. To detect the mCherry, a 561nm (yellow) laser was used with a 616/23 filter and G616 detector. Clonal lines were screened by PCR (Fwd 5’-GAGCAGAGAGGAGGTTAGCG-3’ and Rev 5’-CAAGCTAGACCTTCCCGAGG-3’) to detect excision of the genomic DNA (317 bp) between the two sgRNA cut sites (Figs. S3B).

Clonal *Mgrn1^-/-^* HM1 mouse embryonic stem cells (mESCs) with both GLI-Venus and OLIG2-mKate reporters were previously created using a dual sgRNA strategy and validated (Pusapati et al., 2018a). Similar to what was done in NIH/3T3 cells, clonal double knockout *Mgrn1^-/-^;Rnf157^-/-^* mESC lines were generated using a dual sgRNA strategy to target *Rnf157* in *Mgrn1^-/-^* mESCs. Briefly, the same sgRNAs used to target *Rnf157* in NIH/3T3 cells were used in mESCs, but these sgRNAs were cloned into pSpCas9(BB)-2A-Puro (Addgene) (Ran et al., 2013). Prior to any manipulation, the mESCs were maintained for three passages under feeder free conditions in 2i Medium which contains: DMEM/F12 (Thermo Fisher Scientific, Gibco) and Neurobasal medium (Thermo Fisher Scientific, Gibco) (1:1 ratio) supplemented with 1x N-2 supplement (Thermo Fisher Scientific, Gibco), 1x B-27 supplement (Thermo Fisher Scientific, Gibco), 1% penicillin/streptomycin (Gemini Bioproducts), 2 mM L-Glutamine (Gemini Bioproducts), 55 µM 2-mercaptoethanol (Thermo Fisher Scientific, Gibco), 40 µg/ml bovine serum albumin (MilliporeSigma), 5 µM CHIR 99021 (Axon Medchem), 1 µM PD 98059 (Axon Medchem), and 1000 U/ml ESGRO LIF (MilliporeSigma). Cells were trypsinized in 0.25% Trypsin/EDTA (Thermo Fisher Scientific, Gibco) and rinsed once in PBS. Plasmids were nucleofected into the mESCs using the Lonza Cell nucleofector kit (VAPH-1001) and program A-023 on the Lonza Nucleofector 2b Device (Lonza Bioscience). After the cells were nucleofected, they were plated in 2i Medium onto a 10 cm gelatin-coated CellBIND plate. 24 hours post nucleofection, selection was started and the medium was changed to 2i Medium containing 1.5 µg/ml puromycin (MilliporeSigma) for 48 hours (or until all the cells on the non-nucleofected control plate died). Approximately 1 week after nucleofection, individual mESC colonies were manually picked, expanded, and screened by PCR using the same primers used to screen the *Mgrn1^-/-^;Rnf157^-/-^* NIH/3T3 cells (Fig. S3B).

### Generation of stable cell lines expressing transgenes

Clonal *Megf8^-/-^* and *Mgrn1^-/-^* Flp-In-3T3 cell lines were previously generated and validated (Pusapati et al., 2018a). Stable addback cell lines expressing tagged MEGF8 and MEGF8 ^ΔCtail^ (featured in Figs. 2D and 2E), were generated using Flp recombinase-mediated DNA recombination (Thermo Fisher Scientific, Invitrogen) as previously described (Pusapati et al., 2014). Briefly, the pOG44 Flp-recombinase expression vector (Thermo Fisher Scientific, Invitrogen) and either pEF5/FRT/V5-DEST-*MEGF8*-1D4 or pEF5/FRT/V5-DEST-*MEGF8* ^ΔCtail^-1D4 were transfected into *Megf8^-/-^* NIH/3T3 cells using the X-tremeGENE 9 DNA transfection reagent (Roche Molecular Systems). Approximately 48 hours post transfection the cells were split to 25% confluence and 12-16 hours post split the medium was changed to Complete Medium containing 200 µg/ml Hygromycin B (VWR Life Science). The medium was replenished every 3-4 days and antibiotic selection was conducted for about 2 weeks or until all the cells on the control plate were dead.

Stable addback cell lines expressing tagged MGRN1 (featured in Figs. 2F and 2G) in *Mgrn1^-/-^*; *Rnf157^-/-^* NIH/3T3 cells or tagged MEGF8 in *Megf8^-/-^* NIH/3T3 cells (featured in Figs. 4C and 4D) were generated using the lentiviral expression system. Briefly, to generate lentivirus, four million HEK293T cells were seeded onto a 10 cm plate and 24 hours later these cells were transfected with 1 µg pMD2.G (Addgene), 5 µg psPAX2 (Addgene), and 6 µg of the desired pLenti CMV Puro DEST construct using 36 µl of 1mg/ml polyethylenimine (PEI) (Polysciences). Approximately 48 hours post transfection, the lentivirus was harvested and filtered through a 0.45 µm filter. 2 ml of the filtered lentivirus solution was mixed with 2 ml of Complete Medium containing 16 µg/mL polybrene (MilliporeSigma). The diluted virus was then added to NIH/3T3 cells seeded on 6-well plates. Approximately 48 hours post infection, cells were split and selected with puromycin (2 µg/ml) for 5-7 days or until all the cells on the control plate were dead.

### Established mouse lines

All mouse studies were conducted using animal study protocols approved by the Institutional Animal Care and Use Committee (IACUC) of Stanford University, the University of Pittsburgh, and the McLaughlin Research Institute for Biomedical Sciences. *Mgrn1^md-nc/md-nc^* null mutant mice (referred to in the paper as *Mgrn1^m/m^*) (MGI:3704004) and *Megf8^C193R/C193R^* mice (referred to in the paper as *Megf8^m/m^*) (MGI:3722325) have been described previously (Gunn et al., 2013b; He et al., 2003; Zhang et al., 2009). *Mgrn1^md-nc/+^* animals crossed to *Megf8^C193R/+^* heterozygotes had been outcrossed to FVB/N/Mri and intercrossed for up to 3 generations.

Animals were genotyped for the *Mgrn1^md-nc^* mutation by allele-specific PCR using the following primers: wild-type (Fwd 5’-GCCTGCATGGATAGATGGAT-3’ and Rev 5’-AGGAAGTTGCCCACAAGAACGCA-3’) and mutant (Fwd 5’-CAAGAACAACCAGGAGACTAAGGA-3’ and Rev 5’-GCCCAAGTCCTAAACCTCT-3’).

Amplification was performed using GoTaq Green Master Mix (Promega Corporation), the initial 10 cycles with an annealing temperature of 60 °C, followed by 30 cycles with an annealing temperature of °57 C. Animals were genotyped for the *Megf8^C193R^* mutation by either (1) sequencing of a PCR product generated using primers Fwd 5’-ACGACCCATATCTCTGCCTT-3’ and Rev 5’-GCCTCCAGACCCTCCAAG-3’ or (2) using allele-specific PCR with primers Fwd 5’-CTCAGCTCTGCACCCCTAAC-3’ and Rev (wild-type) 5’-TCCCAAGAATCCAGGTTCACA-3’ or Rev (mutant) 5’-CCAAGAATCCAGGTTCACG-3’. Amplification was performed using GoTaq Green Master Mix (Promega Corporation), 30 cycles with an annealing temperature of 62 °C.

### Generation and validation of *Rnf157^-/-^* mutant mice

*Rnf157^-/-^* mutant mice (referred to in the paper as *Rnf157^m/m^* mice) were generated by CRISPR/Cas9 mediated genome editing. The website Benchling (www.benchling.com) was used to design sgRNAs that target exon 4 of *Rnf157*: (5’-CTACTACCAGGCCACTG-3’ and 5’-TGAACTCGACATTGTAG-3’) (Fig. S3B). Synthetic sgRNAs and Cas9 2NLS nuclease were purchased from Synthego and electroporated into one cell mouse embryos following the Easy Electropopration of Zygotes (EEZy) protocol (Tröder et al., 2018). Briefly, fertilized eggs/1-cell embryos were collected from superovulated C57BL/6J females mated to C57BL/6J males into M2 or EmbryoMax Advanced KSOM medium (MilliporeSigma). Cas9/sgRNA ribonucleoproteins (RNPs) were assembled by combining 4 mM Cas9 protein with 4 mM of sgRNAs in 20 ml Opti-MEM reduced serum medium (ThermoFisher Scientific, Gibco) and incubating 10 min at room temperature. For each electroporation, up to 60 embryos were washed through one drop of Opti-MEM and added to the 20 ml of Cas9 RNP mix. The entire solution was immediately transferred to a 1 mm cuvette (Bio-Rad Laboratories) and placed in a Bio-Rad Gene Pulser XCell electroporator. Two square wave pulses were applied (30V, 3 ms pulse duration, 100 ms interval). Embryos were retrieved from the cuvette by flushing twice with 100 ml of pre-warmed KSOM, transferred to a droplet of KSOM under oil and maintained in a 37 °C incubator with 15% CO_2_ for 1-24 hours. Embryos were subsequently moved through a droplet of M2 medium and transferred to the oviduct of 0.5 dpc (days post coitum) pseudopregnant ICR females. At weaning, a small piece of tail tissue was taken from each pup and the DNA was isolated and genotyped using the following primers to PCR amplify the region around the sgRNA target sequences: Fwd 5’-AACAAAGTCCCGATCCACTG-3’ and Rev1 5’-CAAGCTAGACCTTCCCGAGG-3’ or Rev2 5’-CCTTTCAGCATGGCTTTCTC-3’. Sequence data was analyzed using Synthego’s ICE tool (www.ice.synthego.com/#/) and animals carrying modified alleles predicted to result in a loss of RNF157 function were mated to C57BL/6J animals. *Rnf157^em1Tmg^* carries a single nucleotide deletion at each sgRNA target site: a cysteine at position 58 of exon 4 and another cysteine at position 120 (Fig. S3B). Animals carrying this allele were genotyped by sequencing, as described above, or by allele-specific PCR using the following genotyping primers: Fwd (wild-type) 5’-AGGCAAAGCTAAGGTCCACTAC-3’, Fwd (mutant) 5’-AGGCAAAGCTAAGGTCCACTAA-3’, and Rev 5’-CCTGCTATGCCGTCTTACCT-3’).

RT-PCR was used to verify loss of *Rnf157* expression in *Rnf157^em1Tmg^* mice. Briefly, brains (which express high levels of Rnf157) were collected from wild-type mice and *Rnf157^em1Tmg^* heterozygote and homozygote animals (Fig. S3B). DNase-I treated RNA was extracted using TRIzol reagent (Thermo Fisher Scientific, Invitrogen) and the Direct-zol RNA miniprep kit (Zymo Research). Equal amounts of RNA were used as template for cDNA synthesis using the SuperScript III First-Strand Synthesis System (Thermo Fisher Scientific, Invitrogen). PCR was performed using GoTaq Green Master Mix (Promega Corporation) the following RT-PCR primers: Fwd 5’-ATCCCGTCCAATTCCGTGTA-3’ and Rev 5’-GTACCAGGTGCGATGTAGGA-3’.

## METHOD DETAILS

### Constructs

*MEGF8* constructs: Mammalian Gene Collection (MGC) cDNA clone for human *MEGF8* (NM_001410.3) was purchased from Transomic Technologies, Inc and used as a template for the generation of all *MEGF8* constructs. All *MEGF8* constructs were tagged with a C-terminal 1D4 and cloned into pEF5/FRT/V5-DEST (Thermo Fisher Scientific, Invitrogen) or pLenti CMV PURO DEST (Addgene) using Gateway recombination methods (Thermo Fisher Scientific, Invitrogen). MEGF8^ΔN^ (a.a. 2573-2778) was generated using restriction enzymes SrfI and SapI (New England Biolabs) to remove the N-terminal region (a.a. 26-2572 deleted). MEGF8^ΔCtail^ (a.a. 1-2607) and MEGF8^ΔMASRPFA^ (a.a. 2625-2631 deleted) were created using a combination of overlap extension PCR and restriction enzyme cloning methods. *MEGF8* chimeras were generated using Gibson assembly methods (New England Biolabs). CD16^ECD^-CD7^TM^-MEGF8^Ctail^ (a.a. 2604-2778), CD16^ECD^-MEGF8^TM+Ctail^ (a.a. 2573-2778), and CD16^ECD^-MEGF8^TM+CtailΔMASRPFA^(a.a. 2573-2778 with a.a. 2625-2631 deleted) were all cloned using MEGF8^ΔN^ and CD16^ECD^-CD7^TM^-mCherry-Nck-HA (gift from Bruce Mayer (Rivera et al., 2009)). Lastly, for the bacterial production of MEGF8 Ctail recombinant protein, the C-terminal end of the Ctail (a.a. 2738-2778) was cloned into the pGEX vector using restriction enzyme cloning methods.

*Mgrn1* constructs: Mouse full-length *Mgrn1* (NM_001252437.1) with a C-terminal 3xFLAG tag was synthesized as a gBlock (Integrated DNA Technologies) and used as a template for the generation of all *Mgrn1* constructs. Overlap extension PCR was used to generate MGRN1^Mut1^ (C279A;C282A) and MGRN1^Mut2^ (L307A;R308A). All constructs were cloned into pEF5/FRT/V5-DEST (Thermo Fisher Scientific, Invitrogen) or pLenti CMV PURO DEST (Addgene) using Gateway recombination cloning methods (Thermo Fisher Scientific, Invitrogen).

*Smo* constructs: *mSmo*-EGFP was a gift from Philip Ingham (Zhao et al., 2016). For **Fig. S5C**, pCS2-*mSmo* (Byrne et al., 2016) and a *mSmo* gBlock fragment with all 21 intracellular lysines mutated to arginines (Twist Bioscience) were used as templates to generate the following constructs: untagged full length *Smo* (WT), intracellular lysine-less *Smo* (K0), C-tail lysine-less *Smo* (Ctail^K0^), intracellular loop 2 and 3 lysine-less *Smo* (ICL^K0^), intracellular loop 2 lysine-less *Smo* (ICL2^K0^), and intracellular loop 3 lysine-less *Smo* (ICL3^K0^). Constructs were generated using PCR amplification followed by Gibson assembly methods (New England Biolabs).

Other constructs: SSTR3-GFP was a gift from Kirk Mykytyn (Addgene). pRK5-HA-Ubiquitin-WT and pRK5-HA-Ubiquitin-K0 were purchased from Addgene.

### Reagents and antibodies

Recombinant SHH was expressed in bacteria and purified in the lab as previously described (Bishop et al., 2009). SAG was purchased from Thermo Fisher Scientific (Enzo Life Sciences). The selection antibiotic puromycin was purchased from MilliporeSigma and hygromycin B from VWR Life Science. The transfection reagent XtremeGENE 9 was purchased from Roche Molecular Systems and polybrene from MilliporeSigma. Bafilomycin A1 was purchased from Cayman Chemical and bortezomib was purchased from LC labs. The following primary antibodies were purchased from the following vendors: mouse anti-1D4 (The University of British Columbia, 1:5000); mouse anti-CD16 (clone 3G8, Santa Cruz Biotechnology, 1 µg per 1 million cells in 100ul); mouse anti-CD16 (clone DJ130c, Santa Cruz Biotechnology, 1 µg per 1 million cells in 100ul); mouse anti-FLAG (clone M2, MilliporeSigma, 1:2000); goat anti-GFP (Rockland Immunochemicals, 1:1000); rabbit anti-GFP (Novus Biologicals, 1:5000); mouse anti-GLI1 (clone L42B10, Cell Signaling, 1:1000); mouse anti-HA.11 (clone 16B12, BioLegend, 1:2000); mouse anti-HA (clone 2-2.2.14, Thermo Fisher Scientific, 1:2000); rabbit anti-p38 (Abcam, 1:2000); and rabbit anti-RNF156 (anti-MGRN1, Proteintech, 1:500); mouse anti-ɑ-Tubulin (Clone DM1A, MilliporeSigma, 1:10000); mouse anti-acetylated-Tubulin (MilliporeSigma, 1:10000). The following primary antibodies were generated in the lab or received as a gift: Guinea pig anti-ARL13B (1:1000) (Dorn et al., 2012); rabbit anti-SMO (designed against an intracellular epitope, 1:2000) (Rohatgi et al., 2007b); and rabbit anti-SMO-N (designed against an extracellular epitope, 1:2000) (Milenkovic et al., 2009). The anti-MEGF8 rabbit polyclonal antibody was produced against amino acids 2738-2778 of the mouse MEGF8 protein and affinity purified before use (Cocalico Biologicals, Inc., 1:2000). Hoechst 33342 and secondary antibodies conjugated to horseradish peroxidase (HRP) or Alexa Fluor dyes were obtained from Jackson Laboratories and Thermo Fisher Scientific.

### Protein Sequence Analysis

Iterative sequence profile searches were performed using the PSI-BLAST program run against the NCBI non-redundant (NR) protein database (Altschul et al., 1997). Multiple sequence alignments were built using the Kalign2 software (Lassmann et al., 2009) and were later manually adjusted based on profile-profile, secondary structure information, and structural alignments. Similarity-based clustering for both classification and discarding of nearly identical sequences was performed using the BLASTClust program (Fig. S4A). Maximum-likelihood (ML) tree topology was derived using an edge-linked partition model as implemented in the IQ-TREE software (Nguyen et al., 2015). ModelFinder (Kalyaanamoorthy et al., 2017) was used to automatically identify the best-fit substitution model and estimated “JTT+F+R9” as the suitable model for the given dataset. Branch supports were obtained using the ultrafast bootstrap (UFBoot) approximation method (1000 replicates) (Hoang et al., 2018). To further assess the branch supports, Shimodaira-Hasegawa(SH-)aLRT branch test was also computed as implemented in the IQ-TREE software (Fig. 1B). The sequence logo was generated using the Logo software (Crooks et al., 2004) (Fig. 2B). An alignment comprising a collection of all unique members of the MEGF8-Attractin family from the RefSeq database was utilized as input. The UniProt align tool was used to compare two protein sequences with the Clustal Omega program (Fig. S3A) (UniProt Consortium, 2019). Sequence analysis of the MGRN1 RING domain was done using ConSurf (Ashkenazy et al., 2016). Briefly, 200 MGRN1 homologs were collected from UniProt using the homolog search algorithm HMMER and a color coded multiple sequence alignment was built using ClustalW (Fig. S4B).

### Immunoprecipitation and Western Blotting

Whole cell extracts from HEK293T and NIH/3T3 cells were prepared in Immunoprecipitation (IP) Lysis Buffer containing: 50 mM Tris at pH 8.0, 150 mM NaCl, 1% NP-40, 1 mM DTT, 1x SIGMAFAST protease inhibitor cocktail (MilliporeSigma), and 1x PhosSTOP phosphatase inhibitor cocktail (Roche). Cells were lysed for 1 hour on a shaker at 4 °C, supernatants were clarified by centrifugation, and 1D4 tagged MEGF8 was captured by a 1D4 antibody (The University of British Columbia) covalently conjugated to Protein A Dynabeads (Thermo Fisher Scientific, Invitrogen). Immunoprecipitates were washed once with IP Wash Buffer A (50 mM Tris at pH 8.0, 150 mM NaCl, 1% NP-40, and 1 mM DTT), once with wash IP Wash Buffer B (50 mM Tris at pH 8.0, 500 mM NaCl, 0.1% NP-40, and 1 mM DTT), and finally with IP Wash Buffer C (50 mM Tris at pH 8.0, 0.1% NP-40, and 1 mM DTT). Proteins were eluted by resuspending samples in 1xNuPAGE LDS sample buffer (Thermo Fisher Scientific, Invitrogen) supplemented with 100 mM DTT, incubated at 37 °C for 30 min, and subjected to SDS-PAGE (Figs. 2C, 2D, and S4C).

For all other immunoblotting data presented in the manuscript, whole cell extracts were prepared in RIPA lysis buffer containing: 50 mM Tris at pH 8.0, 150 mM NaCl, 2% NP-40, 0.25% Deoxycholate, 0.1% SDS, 0.5 mM TCEP, 10% glycerol, 1x SIGMAFAST protease inhibitor cocktail (MilliporeSigma), and 1x PhosSTOP phosphatase inhibitor cocktail (Roche). The resolved proteins were transferred onto a nitrocellulose membrane (Bio-Rad Laboratories) using a wet electroblotting system (Bio-Rad Laboratories) followed by immunoblotting.

### Flow cytometry of live cells

As described above, a lentiviral expression system was used to stably express CD16/CD7/MEGF8 chimeras in *Megf8^-/-^* cells NIH/3T3 cells (diagramed in Fig. S5D). A modified live cell immunostaining protocol from Santa Cruz Biotechnology and Cell Signaling Technology was used to label and analyze cell surface CD16/CD7/MEGF8 chimeras (Fig. S5E). Briefly, four cell lines were analyzed: *Megf8^-/-^*, *Megf8^-/-^* with *CD16^ECD^-CD7^TM^-Megf8^Ctail^* addback, *Megf8^-/-^* with *CD16^ECD^-Megf8^TM+Ctail^* addback, and *Megf8^-/-^* with *CD16^ECD^-Megf8^TM+CtailΔMASRPFA^* addback. Prior to staining, the cells were serum starved for 24 hours to allow for primary cilia growth. On staining day, the growth medium was removed, the cells were rinsed with PBS, and then dissociated in 0.2% EDTA (prepared in PBS) for approximately 5 minutes at 37°C. Upon seeing the cells lift from the plate, the cells were pipetted up and down five times to create a single cell suspension, and Complete Medium (containing 10% FBS) was added to neutralize the EDTA. A small sample was taken to determine the total number of cells present. The cells were then resuspended in Flow Cytometry (FCM) Blocking Buffer (0.5% bovine serum albumin prepared in PBS) at a concentration of 10 million cells/ml. The cells were blocked for 10 min on ice, 1 million cells (100 ul of the cell suspension) was then transferred to a fresh tube, and 1 ug of an anti-CD16 antibody (Santa Cruz Biotechnology, clones 3G8 and DJ130c) was added directly to the cells. Primary antibodies were administered for 30 minutes on ice. Primary antibodies were rinsed off with 2 washes in FCM Blocking Buffer. The cells were then incubated for 30 minutes on ice in 1 ug of donkey anti-mouse IgG, Alexa Fluor 488 (Thermo Fisher Scientific, Invitrogen) diluted in FCM Blocking Buffer. The cells were washed 2 times in FCM Blocking Buffer then analyzed on a BD Accuri C6 Flow Cytometer (BD Biosciences).

### SMO internalization assay

Cell surface internalization assay for SMO was performed as described previously for Fig. 3 (Pusapati et al., 2018a). Briefly, wild type, *Megf8^-/-^*, and *Mgrn1^-/-^*;*Rnf157^-/-^* NIH/3T3 cells were plated on 15 cm plates in Complete Medium (containing 10% FBS). Once the cells were confluent they were switched to Low Serum Medium (complete medium containing 0.5% FBS) for 24 hours. On biotinylation day, the cells were removed from the 37 °C incubator and placed on an ice-chilled metal rack in a 4 °C cold room. The medium was removed and cells were quickly washed 3 times with ice-cold DPBS+ buffer (Dulbecco’s PBS supplemented with 0.9 mM CaCl_2_, 0.49 mM MgCl_2_.6H_2_O, 5.6 mM dextrose, and 0.3 mM sodium pyruvate). Biotinylation of cell surface proteins using a non-cell permeable and thiol-cleavable probe was initiated by incubating cells with 0.4 mM EZ-Link Sulfo-NHS-SS-Biotin (Thermo Fisher Scientific) in DPBS+ buffer for 30 minutes. Unreacted Sulfo-NHS-SS-Biotin was quenched with 50 mM Tris (pH 7.4) for 10 min. Cells were then washed 3 times with a 1x Tris-buffered saline (25 mM Tris at pH 7.4, 137 mM NaCl, and 2.7 mM KCl) and whole cell extracts were prepared in Biotinylation Lysis Buffer A (50 mM Tris at pH 8.0, 150 mM NaCl, 2% NP-40, 0.25% Deoxycholate, 1x SIGMAFAST protease inhibitor cocktail (MilliporeSigma), and 1x PhosSTOP phosphatase inhibitor cocktail (Roche). Biotinylated proteins from clarified supernatants were captured on a streptavidin agarose resin (TriLink Biotechnologies), washed once with Biotinylation Lysis Buffer A, once with Biotinylation Wash Buffer A (Biotinylation Lysis Buffer A + 0.5% SDS), once with Biotinylation Wash Buffer B (Biotinylation Wash Buffer A + 150 mM NaCl), and finally once again with Biotinylation Wash Buffer A. Biotinylated proteins captured on streptavidin agarose resin were eluted in 1x NuPAGE-LDS sample buffer (Thermo Fisher Scientific, Invitrogen) containing 100 mM DTT at 37 °C for 1 hour and assayed by immunoblotting for SMO (Figs. 3B and 3C).

### Ubiquitination assay

8 million HEK293T cells were plated onto a 15 cm plate. 24 hours after plating, the cells were transfected using PEI. 6 ug of each construct was transfected into the cells (at a DNA:PEI ratio of 1:3). An empty plasmid construct was used as filler DNA to ensure that each plate was transfected with the same amount of DNA. 36 hours post transfection, cells were pre-treated with 10 µM bortezomib (a proteasome inhibitor) and 100 nM bafilomycin A1 (a lysosome inhibitor) for 4 hours to enrich for ubiquitinated proteins. Cells were washed twice with chilled 1x PBS and lysed in Ubiquitination Lysis Buffer A comprised of: 50 mM Tris at pH 8.0, 150 mM NaCl, 2% NP-40, 0.25% sodium deoxycholate, 0.1% SDS, 6M urea, 1 mM DTT, 10 µM bortezomib, 100 nM bafilomycin A1, 20 mM N-Ethylmaleimide (NEM, MilliporeSigma), and 1x SIGMAFAST protease inhibitor cocktail (MilliporeSigma). Clarified supernatants were diluted ten-fold with Ubiquitination Lysis Buffer B (Ubiquitination Lysis Buffer A prepared without urea) to adjust the urea concentration to 600 mM. For these assays, we assessed ubiquitination on both GFP tagged and untagged SMO. Ubiquitinated GFP tagged SMO (Figs. 4A, 4B, S5A, and S5D) was captured using a GFP binding protein (GBP) covalently conjugated to carboxylic acid decorated Dynabeads (Dynabeads M-270 carboxylic acid, Thermo Fisher Scientific). Untagged SMO (Fig. S5C) was captured using SMO antibody covalently conjugated to Protein A Dynabeads (Thermo Fisher Scientific, Invitrogen). Immunoprecipitates were washed once with Ubiquitination Wash Buffer A (Ubiquitination Lysis Buffer B + 0.5% SDS), once with Ubiquitination Wash Buffer B (Ubiquitination Wash Buffer A + 1 M NaCl), and finally once again with Ubiquitination Wash Buffer A. Proteins bound to dynabeads were eluted in 2x NuPAGE-LDS sample buffer (Thermo Fisher Scientific, Invitrogen) containing 30 mM DTT at 37 °C for 30 minutes and assayed by immunoblotting for GFP or SMO antibodies for GFP tagged SMO and endogenous SMO, respectively.

### Immunofluorescence staining of cells and tissue and image quantifications

Mouse embryos (e12.5) were harvested and fixed in 4% (w/v) paraformaldehyde (PFA) in 1x PBS for 2 hours at 4 °C and then rinsed thoroughly in chilled PBS. To cryopreserve the tissue, the embryos were transferred to 30% sucrose in 0.1M PB (pH 7.2) and allowed to equilibrate overnight. To allow for better analysis of the tissue, the embryos were further dissected into five pieces: 2 hands (forelimbs), head, upper body, and lower body. All five pieces were then mounted and frozen into Tissue-Plus OCT (optimal cutting temperature) compound (Thermo Fisher Scientific) and 12-14 µm sections were collected. Prior to staining, the tissue was blocked for 1 hour in immunofluorescence (IF) Blocking Buffer containing: 1% normal donkey serum (NDS) and 0.1% Triton-X diluted in 1x PBS. In a humidified chamber, the sections were incubated with primary antibodies overnight at 4 °C, rinsed 3 times in PBST (1x PBS + 0.1% Triton-X), incubated with secondary antibodies and Hoescht for 1 hour at room temperature, rinsed 3 times in PBST, and then mounted in Prolong Gold antifade mountant (Thermo Fisher Scientific, Invitrogen).

NIH/3T3 cells, pMEFs, and primary human fibroblasts were fixed in chilled 4% PFA in 1x PBS for 10 minutes and then rinsed with chilled PBS. Cells were incubated in IF Blocking Buffer for 30 minutes, primary antibodies for 1 hour, and secondary antibodies for 30 minutes.

Fluorescent images were acquired on an inverted Leica SP8 confocal microscope equipped with a 63X oil immersion objective (NA 1.4). Z-stacks (∼4 µm sections) were acquired with identical acquisition settings (laser power, gain, offset, frame and image format) within a given experiment. An 4-8X optical zoom was used for imaging cilia to depict representative images. For the quantification of SMO at cilia, images were opened in Fiji with projections of the maximum fluorescent intensities of z-stacks. Ciliary masks were constructed based on ARL13B images and then applied to corresponding SMO images to measure the fluorescence intensity of SMO at cilia.

### Mouse embryo phenotyping analysis

Mouse embryos (e13.5-14.5) were fixed in 4% (w/v) PFA in 1x PBS for 2-3 days. Necropsy was performed to determine visceral organ situs (i.e. lung and liver lobation, heart and stomach situs, and spleen and pancreas structure). The samples were embedded in paraffin and processed for episcopic confocal microscopy as previously described (Liu et al., 2013). Briefly, this entailed sectioning of the tissue block using a Leica sledge microtome with serial images of the block face captured with a Leica confocal microscope. The serial two-dimensional (2D) image stacks generated were three-dimensionally (3D) reconstructed using the Osirix software (Rosset et al., 2004) and digitally resliced in different orientations to aid in the analysis of intracardiac anatomy and the diagnosis of congenital heart defects (Liu et al., 2013).

### Variant Discovery and Validation

Genomic DNA was extracted from blood using the PAXgene Blood DNA kit (Qiagen). Patient genomic DNA was analyzed using whole exome sequencing performed using the Agilent V5 Exome Capture kit followed by sequencing with the Illumina HiSeq2000 with 150 base paired-end reads with 100X coverage. Reads were aligned to the human reference genome (version hg19) using Burrows-Wheeler Alignment (BWA, version 0.5.9) (Li and Durbin, 2009) with default parameters, and further processed according to the recommendations of the Genome Analysis Toolkit (GATK) Best Practices (Auwera and Others, 2016; DePristo et al., 2011). GATK HaplotypeCaller was used for single-nucleotide polymorphism (SNP) and insertion/deletion mutation (INDEL) discovery and variants that passed the GATK Variant Score Quality Recalibration (VQSR) and standard GATK filters with minor allele frequency <5% based on the Genome Aggregation Database (GnomAD). Only variants with Combined Annotation-Dependent Depletion (CADD) PHRED (Kircher et al., 2014) score of at least 10 were considered, and PolyPhen-2 (Adzhubei et al. 2010) and SIFT (Kumar, Henikoff, and Ng 2009) were used to assess variant pathogenicity. MEGF8/MGRN1/RNF157 variants recovered were validated by Sanger sequencing and heritable transmission was determined by further Sanger sequencing of genomic DNA from the parents.

## QUANTIFICATION AND STATISTICAL ANALYSIS

All data analysis and graphs were generated using GraphPad Prism 8. Violin plots were created using the “Violin Plot (truncated)” appearance function. In Prism 8, the frequency distribution curves of the violin plots are calculated using kernal density estimation. By using the “truncated” violin plot function, the frequency distributions shown are confined within the minimum to maximum values of the data set. On each violin plot, the median (central bold line) and quartiles (adjacent thin lines, representing the first and third quartiles) are labeled.

In Prism 8, all statistical tests were conducted using non-parametric methods, which do not assume a Gaussian distribution of the data. The statistical significance between two groups was determined using the Mann-Whitney test and the significance between three or more groups was determined using the Kruskal-Wallis test. For each figure, the error bars (representing the standard deviation) and *p*-values were all calculated using Prism 8 and reported in the figure legend. *P*-values were reported using the following key: not-significant (ns) *p*-value > 0.05, \**p*-value ≤ 0.05, \*\**p*-value ≤ 0.01, \*\*\**p*-value ≤ 0.001, and \*\*\*\**p*-value ≤ 0.0001 Additional figure details regarding the *n*-value and statistical test applied were also reported in the individual figure legends.

All cell biological and biochemical experiments were performed two to three independent times, with similar results. To validate newly generated *Mgrn1^-/-^*;*Rnf157^-/-^* NIH/3T3 and neural progenitor cell lines, 3 independent clonal cell lines were analyzed. Analysis of one clonal NIH/3T3 cell line was featured in the main figures (Figs. 1A and 1C) and data from the additional cell lines was presented in Figs. S3C, S3D, and S3E. Similarly, 2-3 primary mouse embryonic fibroblast (pMEF) cell lines were analyzed from both *Megf8^m/m^* and *Mgrn1^m/m^* embryos, where each pMEF cell line was generated from a single embryo (Figs. S1B and S1C).

## DATA AND CODE AVAILABILITY

The published article contains all datasets generated and analyzed during this study.

## File S1: Newick tree file for the MGRN1-RNF157 family, Related to Figure 1B

(MGRN1_XP_626702.1_Cryptosporidium_parvum:6.4556799939,((((MGRN1_XP_012750170.1_Acytostelium_subglobosum:0.8834035749,MGRN1_EFA84789.1_Heterostelium_album:0.756 4698726)100:1.0990476127,MGRN1_XP_646206.1_Dictyostelium_discoideum:1.9725849426) 100:3.1996779932,((((((((((((((MGRN1_XP_022252119.1_Limulus_polyphemus:0.7130424361, MGRN1_XP_002434259.1_Ixodes_scapularis:0.8067633411)100:0.3546045995,(((MGRN1_X P_021208538.1_Bombyx_mori:0.0000022207,MGRN1_XP_012551644.1_Bombyx_mori:0.000 0022207)100:0.6674373775,(MGRN1_XP_008197724.1_Tribolium_castaneum:0.6043109157,((MGRN1_XP_624563.2_Apis_mellifera:0.0000022207,MGRN1_XP_006571957.1_Apis_mellifer a:0.0000022207)100:0.0978541753,MGRN1_XP_011144180.1_Harpegnathos_saltator:0.1639 699560)100:0.4188667910)100:0.1383218127)97:0.1390462809,(MGRN1_NP_572915.1_Dros ophila_melanogaster:0.7240618615,MGRN1_XP_001842757.1_Culex_quinquefasciatus:0.493 0549513)100:0.5592043648)98:0.1659530383)99:0.1601768095,MGRN1_XP_003378072.1_Tr ichinella_spiralis:2.4613076368)96:0.1652080042,MGRN1_OWA53600.1_Hypsibius_dujardini: 1.8845700706)100:0.3022685831,((((MGRN1_XP_009012902.1_Helobdella_robusta:1.419935 0327,MGRN1_XP_011456291.1_Crassostrea_gigas:0.9744293918)68:0.1827551179,MGRN1_ XP_013395504.1_Lingula_anatina:0.9435380355)68:0.1936942442,MGRN1_XP_014772328.1_Octopus_bimaculoides:0.8240707803)77:0.2250694468,MGRN1_XP_009056095.1_Lottia_gig antea:0.9606406571)100:0.3497208259)98:0.2460667652,((((MGRN1_XP_015752852.1_Acropora_digitifera:1.8098585308,MGRN1_KXJ08562.1_Exaiptasia_pallida:0.2865159553)100:0.89 20209580,MGRN1_XP_001630091.1_Nematostella_vectensis:0.6942822428)100:1.1606397688,(MGRN1_XP_002109227.1_Trichoplax_adhaerens:2.2929217319,MGRN1_XP_012558349.1_Hydra_vulgaris:2.6718292265)95:0.4159482970)94:0.2982637166,MGRN1_XP_018671577.1_Ciona_intestinalis:2.4449979012)45:0.1600650448)40:0.1743552262,(MGRN1_XP_011681930.1_Strongylocentrotus_purpuratus:1.2412630478,MGRN1_XP_006816719.1_Saccoglossus_k owalevskii:0.6956647937)100:0.3512516240)39:0.1272151383,MGRN1_XP_002607160.1_Branchiostoma_floridae:1.5874372991)86:0.4011963809,((((MGRN1_NP_001138254.1_Danio_reri o:0.1542059135,MGRN1_CAG12208.1_Tetraodon_nigroviridis:0.0900101225)100:0.09655401 88,MGRN1_XP_017339726.1_Ictalurus_punctatus:0.5529251973)100:0.1053269429,(MGRN1_XP_018096102.1_Xenopus_laevis:0.6339067055,(((MGRN1_XP_006521895.1_Mus_musculu s:0.0881653705,MGRN1_ELK34685.1_Myotis_davidii:0.2355299819)99:0.0271033339,MGRN 1_NP_056061.1_Homo_sapiens:0.1092290169)100:0.1239586071,MGRN1_XP_004945431.2_ Gallus_gallus:0.1734568780)99:0.0546549251)98:0.0676536127)100:0.7446594670,(((((((((((RNF157_XP_019794160.1_Tursiops_truncatus:0.0715318657,(RNF157_XP_007454376.1_Lipot es_vexillifer:0.0078127635,RNF157_XP_012390593.1_Orcinus_orca:0.0417563415)100:0.003 9693380)100:0.0274949628,RNF157_OWK14388.1_Cervus_elaphus:0.0343731679)96:0.0138497635,(((RNF157_XP_014387228.1_Myotis_brandtii:0.5339584241,(RNF157_XP_015982952.1_Rousettus_aegyptiacus:0.0641276630,(RNF157_ELK12342.1_Pteropus_alecto:0.00214670 44,RNF157_XP_011360372.1_Pteropus_vampyrus:0.2733353942)99:0.0222111173)99:0.0432625583)100:0.0410727678,((((RNF157_XP_006107235.1_Myotis_lucifugus:0.0209344513,RN F157_XP_015414699.1_Myotis_davidii:0.0757533803)100:0.0969409524,RNF157_XP_013845346.1_Sus_scrofa:0.1680616436)88:0.0743225206,(RNF157_XP_004655748.1_Jaculus_jacul us:0.0764856949,RNF157_XP_005624217.1_Canis_lupus:0.0583989552)87:0.0142601111)80:0.0116044543,(RNF157_XP_017529823.1_Manis_javanica:0.0678685327,(RNF157_XP_007536806.1_Erinaceus_europaeus:0.0259028420,(RNF157_XP_012588480.1_Condylura_cristata: 0.0883633390,RNF157_XP_012789771.1_Sorex_araneus:0.1336762544)100:0.0135134547)99:0.0082482797)96:0.0098075317)79:0.0075837417)68:0.0120886129,RNF157_EQB77236.1_ Camelus_ferus:0.2062033966)93:0.0215268641)97:0.0053043244,RNF157_XP_019500002.1_Hipposideros_armiger:0.0425196353)98:0.0082871101,((((RNF157_XP_017391079.1_Cebus_ capucinus:0.0635807445,((RNF157_AAH04231.2_Homo_sapiens:0.3429153408,RNF157_NP_ 443148.1_Homo_sapiens:0.0039774845)99:0.0153998507,(RNF157_XP_017740235.1_Rhinopithecus_bieti:0.0389188257,RNF157_XP_008009863.1_Chlorocebus_sabaeus:0.0699256935) 100:0.0043091587)99:0.0019014587)99:0.0106880921,RNF157_XP_012658433.1_Otolemur_ garnettii:0.0626974246)100:0.0035982813,(RNF157_NP_081534.1_Mus_musculus:0.0648491 574,RNF157_XP_004592943.1_Ochotona_princeps:0.0864653650)99:0.0049199899)100:0.0077707120,RNF157_XP_004374441.1_Trichechus_manatus:0.0820793497)95:0.0036907473)99:0.0419720673,RNF157_XP_016083403.1_Ornithorhynchus_anatinus:0.3883801346)93:0.0121062608,RNF157_XP_003768510.1_Sarcophilus_harrisii:0.0747707403)100:0.0664719130,((((RNF157_XP_014435712.1_Pelodiscus_sinensis:0.0847040242,RNF157_XP_008162104.1_C hrysemys_picta:0.3616339896)95:0.0326348156,RNF157_XP_006018423.2_Alligator_sinensis:0.0482559786)91:0.0347732273,((RNF157_XP_009664550.1_Struthio_camelus:0.057139492 2,RNF157_XP_013797312.1_Apteryx_mantelli:0.0629791352)98:0.0332317156,((RNF157_XP_010299131.1_Balearica_regulorum:0.5279873468,((((RNF157_XP_009914760.1_Haliaeetus_ albicilla:0.1297778575,(((RNF157_XP_021270628.1_Numida_meleagris:0.0158663718,RNF15 7_XP_015150798.1_Gallus_gallus:0.0100129247)100:0.0838907874,RNF157_XP_009562266.1_Cuculus_canorus:0.0499423928)98:0.0155239002,RNF157_XP_009636045.1_Egretta_garz etta:0.1132778761)58:0.0000026051)92:0.0139762340,RNF157_XP_010142984.1_Buceros_rhinoceros:0.0399010091)78:0.0084636664,((((RNF157_XP_012433989.1_Taeniopygia_guttata: 0.3336668594,RNF157_XP_021397362.1_Lonchura_striata:0.0317956558)99:0.0088702849,R NF157_XP_016158280.1_Ficedula_albicollis:0.0848070169)87:0.0126059415,(RNF157_XP_014127545.1_Zonotrichia_albicollis:0.0996621112,(RNF157_XP_019146210.1_Corvus_cornix:0. 0388465764,RNF157_XP_017688572.1_Lepidothrix_coronata:0.0346467421)90:0.0117236358)87:0.0282220494)92:0.0187083207,(RNF157_KQK80228.1_Amazona_aestiva:0.0459980143,RNF157_XP_012983257.1_Melopsittacus_undulatus:0.1876689248)100:0.0931602731)85:0.0 182830049)51:0.0000024413,RNF157_XP_019327654.1_Aptenodytes_forsteri:0.1300043743)72:0.0133699745)74:0.0096468713,(RNF157_XP_008498014.1_Calypte_anna:0.1579187411, RNF157_XP_010193349.1_Mesitornis_unicolor:0.0531898268)56:0.0085821787)99:0.1064356 370)100:0.0904064361)57:0.0126676955,(RNF157_XP_016846820.1_Anolis_carolinensis:0.0743760823,(RNF157_ETE73233.1_Ophiophagus_hannah:0.1534674086,RNF157_XP_0152699 26.1_Gekko_japonicus:0.1622902018)57:0.0190621432)100:0.0680287271)100:0.0513493674)100:0.0602604858,(RNF157_XP_018092914.1_Xenopus_laevis:0.1062829246,RNF157_XP_ 018410793.1_Nanorana_parkeri:0.0548212369)100:0.3383893646)99:0.1080653267,RNF157_ XP_014342274.1_Latimeria_chalumnae:0.2320355122)96:0.0467261359,((((((((RNF157_XP_005952432.2_Haplochromis_burtoni:0.2853462810,(RNF157_XP_008294193.1_Stegastes_parti tus:0.0070064422,((RNF157_XP_020792200.1_Boleophthalmus_pectinirostris:0.2404359276,R NF157_XP_015818331.1_Nothobranchius_furzeri:0.0611955329)98:0.0185998689,RNF157_X P_018521135.1_Lates_calcarifer:0.0122103587)85:0.0057484926)80:0.0000024663)93:0.0101518814,RNF157_XP_003978203.2_Takifugu_rubripes:0.0825142682)100:0.0711936727,((RNF157_XP_021436693.1_Oncorhynchus_mykiss:0.0265075529,RNF157_XP_014068740.1_Salm o_salar:0.0279331129)100:0.1176562479,RNF157_XP_012987491.1_Esox_lucius:0.07848920 20)100:0.0647841386)100:0.0741729306,RNF157_XP_012691288.1_Clupea_harengus:0.2435050180)99:0.0160095459,((RNF157_XP_017558155.1_Pygocentrus_nattereri:0.0285236736,R NF157_XP_017338147.1_Ictalurus_punctatus:0.1652275577)100:0.0864146530,RNF157_XP_ 018963967.1_Cyprinus_carpio:0.0972291116)99:0.0462892325)99:0.0428323245,RNF157_XP_018617615.1_Scleropages_formosus:0.0990364186)94:0.0435787077,RNF157_XP_015211982.1_Lepisosteus_oculatus:0.1570006008)100:0.2679641271,(RNF157_XP_007884001.1_Call orhinchus_milii:0.3257010038,RNF157_XP_020381796.1_Rhincodon_typus:0.5614613586)97: 0.1285779675)93:0.0265497081)100:0.6580846945)100:0.4900158004)99:0.4651716473,MGRN1_XP_019849223.1_Amphimedon_queenslandica:3.6035969064)62:0.1807345037,((MGRN 1_CBY07841.1_Oikopleura_dioica:2.7847834568,MGRN1_OAF67975.1_Intoshia_linei:4.06784 41542)84:0.7054571260,(MGRN1_NP_510385.1_Caenorhabditis_elegans:1.5469957734,(MG RN1_XP_001892209.1_Brugia_malayi:0.5608706112,MGRN1_ERG87446.1_Ascaris_suum:0. 6510690897)100:1.0134141185)100:1.1752088817)56:0.4547444207)95:0.6156688389,((((MGRN1_XP_004346075.1_Capsaspora_owczarzaki:1.2611631116,MGRN1_XP_014158540.1_Sp haeroforma_arctica:3.5625632456)98:0.8367261412,MGRN1_XP_004992715.1_Salpingoeca_r osetta:2.2705221465)94:1.1062203011,(MGRN1_XP_013760930.1_Thecamonas_trahens:4.65 14487514,MGRN1_XP_009496900.1_Fonticula_alba:7.1644480376)70:2.3964454855)58:0.7467699989,(((MGRN1_ORX91900.1_Basidiobolus_meristosporus:2.5042194473,(MGRN1_KNE 56649.1_Allomyces_macrogynus:2.1174716183,MGRN1_ORZ33642.1_Catenaria_anguillulae: 2.1224826073)100:3.6582738324)100:0.8564694622,MGRN1_KFH67492.1_Mortierella_verticillata:3.1705019833)100:1.5530686992,(MGRN1_XP_006675586.1_Batrachochytrium_dendrob atidis:3.3171805662,MGRN1_XP_016604932.1_Spizellomyces_punctatus:2.6740192696)100:1.7578620674)96:0.8431626334)61:0.4916394823)87:0.4672283754,((((((MGRN1_XP_005786359.1_Emiliania_huxleyi:2.7875210880,MGRN1_XP_005819899.1_Guillardia_theta:1.86572626 31)99:1.0565410313,(((((MGRN1_XP_011396820.1_Auxenochlorella_protothecoides:1.419388 2969,MGRN1_XP_005849270.1_Chlorella_variabilis:0.7250397887)100:0.3787123806,MGRN1_XP_005648141.1_Coccomyxa_subellipsoidea:0.9471512839)100:1.1869157110,MGRN1_X P_001703291.1_Chlamydomonas_reinhardtii:1.7371911370)100:0.7223991659,((MGRN1_XP_ 002508870.1_Micromonas_commoda:1.3760598152,MGRN1_XP_003063967.1_Micromonas_ pusilla:1.5889584546)100:0.5886215128,MGRN1_XP_003083824.1_Ostreococcus_tauri:3.568 1305083)100:0.8763305688)99:0.4492440239,(((((MGRN1_XP_015159045.1_Solanum_tuberosum:0.9988391110,MGRN1_NP_566356.1_Arabidopsis_thaliana:0.8353173904)100:0.403917 0893,MGRN1_NP_001333932.1_Zea_mays:0.6619418321)100:0.8913461077,(MGRN1_XP_016464410.1_Nicotiana_tabacum:0.6756830166,MGRN1_XP_010659828.1_Vitis_vinifera:0.776 3921462)100:1.4485243926)99:0.2272140046,(MGRN1_XP_001764144.1_Physcomitrella_patens:1.1831346121,MGRN1_XP_002982812.1_Selaginella_moellendorffii:0.9015411581)83:0.2 312402188)100:0.8165187923,MGRN1_G054.1_Klebsormidium_nitens:1.6139073283)100:0.4675215425)99:0.9331956584)94:0.4040903123,(((MGRN1_XP_012193830.1_Saprolegnia_parasitica:0.1115517198,MGRN1_XP_008607414.1_Saprolegnia_diclina:0.0641701944)100:0.7326918078,MGRN1_XP_009843084.1_Aphanomyces_astaci:1.2869041723)100:1.18332501 69,MGRN1_XP_009529847.1_Phytophthora_sojae:1.2771853540)100:2.7188904758)88:0.5582954625,(((MGRN1_XP_001612895.1_Plasmodium_vivax:3.4315460842,MGRN1_CEM20353.1_Vitrella_brassicaformis:1.3890543205)100:0.6916400651,(MGRN1_XP_003880832.1_Neosp ora_caninum:0.2718422860,(MGRN1_XP_002368812.1_Toxoplasma_gondii:0.0167429888,M GRN1_XP_008887669.1_Hammondia_hammondi:0.0637855327)80:0.1327641691)99:1.76226 95251)99:1.1437754151,MGRN1_XP_002769107.1_Perkinsus_marinus:4.1019957488)99:1.6755472339)76:0.4117119488,MGRN1_EWM25530.1_Nannochloropsis_gaditana:4.8947789704)62:0.4236757775,(MGRN1_XP_005706327.1_Galdieria_sulphuraria:4.1659250600,MGRN1_X P_005536920.1_Cyanidioschyzon_merolae:3.2719392440)92:1.4899541461)54:0.3940419836)48:0.3068887302,(MGRN1_CEO98225.1_Plasmodiophora_brassicae:3.6368224377,MGRN1_ XP_002676307.1_Naegleria_gruberi:4.8727155631)94:1.1899609077)36:0.1822088386)78:0.1936497834,(((MGRN1_XP_001443657.1_Paramecium_tetraurelia:5.0710744593,MGRN1_XP_001025361.3_Tetrahymena_thermophila:4.9804036755)86:0.7319329411,MGRN1_EJY88708. 1_Oxytricha_trifallax:5.1625158756)86:1.1676874607,((((MGRN1_EPY43869.1_Angomonas_d eanei:0.5516396833,(MGRN1_XP_015657042.1_Leptomonas_pyrrhocoris:0.6159047219,MGR N1_CCW69607.1_Phytomonas_sp.:0.4633390492)100:0.1611523546)100:0.6005248731,MGR N1_XP_815262.1_Trypanosoma_cruzi:0.6064438770)100:4.9327798931,(MGRN1_XP_953663.1_Theileria_annulata:5.0422314388,MGRN1_XP_012766930.1_Babesia_bigemina:4.031483 2781)99:2.2203882408)84:0.9561853312,MGRN1_XP_012895226.1_Blastocystis_hominis:6.2068238081)59:0.0855120691)64:0.3231918371)60:0.2626447148,(((MGRN1_XP_847288.1_Trypanosoma_brucei:1.2783452619,MGRN1_XP_009310993.1_Trypanosoma_grayi:1.04965301 44)100:0.7271364166,(MGRN1_EPY22903.1_Strigomonas_culicis:2.7318592994,(MGRN1_KPI86947.1_Leptomonas_seymouri:1.2291044873,MGRN1_XP_001463767.1_Leishmania_infant um:1.1355401363)100:1.2104190489)100:0.6619769589)100:5.7799142426,MGRN1_OMJ90811.1_Stentor_coeruleus:4.0533298055)83:0.9350947891);

---

**Figure S1:**
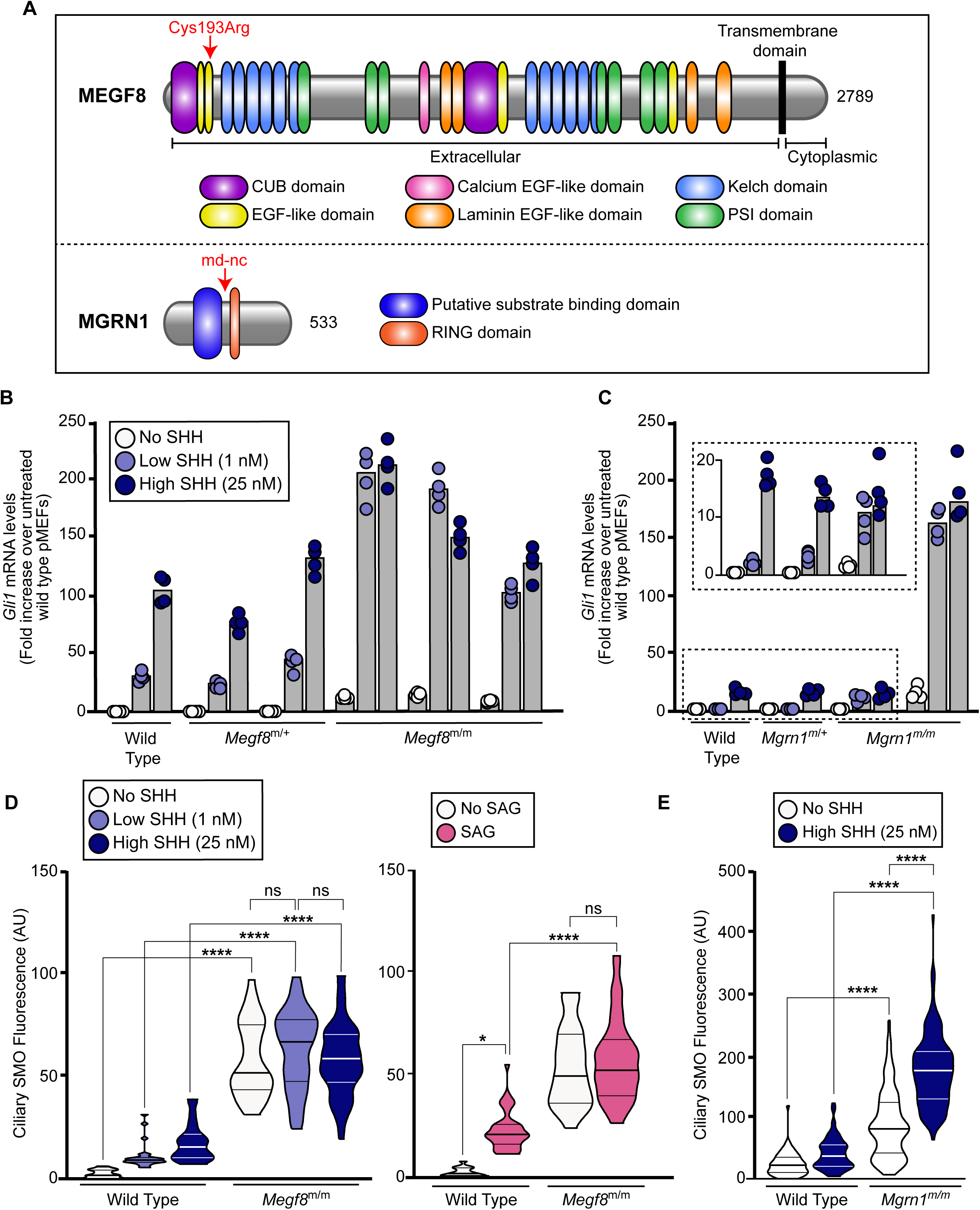
Primary fibroblasts from *Megf8^m/m^* and *Mgrn1^m/m^* mouse embryos display elevated Hh signaling and ciliary SMO abundance, Related to Figure 1. **(A)** Domain composition of mouse MEGF8 and MGRN1 (UniProt Consortium, 2019). The arrow above MEGF8 denotes the location of the missense mutation (C193R) in the second EGF-like domain of MEGF8 (Zhang et al., 2009) that causes heart defects and heterotaxy in mice. In the loss-of-function *md-nc* allele of *Mgrn1*, a thymidine to adenine mutation in intron 9 disrupts splicing, leading to a premature stop (at the position shown by the arrow above MGRN1) and nonsense mediated decay of the transcript (He et al., 2003). **(B and C)** Hh signaling strength was assessed using qRT-PCR to measure *Gli1* mRNA in primary mouse embryonic fibroblasts (pMEFs) prepared from (B) wild-type, *Megf8^m/+^*, and *Megf8^m/m^* embryos or (C) wild-type, *Mgrn1^m/+^*, and *Mgrn1^m/m^* embryos treated with no, low (1 nM), or high (25 nM) concentrations of SHH. Each cell line tested was derived from a different embryo. Bars denote the median *Gli1* mRNA values derived from the four individual measurements shown as circles. **(D and E)** Violin plots summarizing quantification of SMO fluorescence at ∼15-50 cilia from pMEFs of the indicated genotypes treated treated with no Hh agonist, low SHH (1 nM), high SHH (25 nM), or the direct SMO agonist SAG (100 nM). Statistical significance was determined by the Kruskal-Wallis test; not-significant (ns) > 0.05, \**p*-value ≤ 0.05, and \*\*\*\**p*-value ≤ 0.0001.

**Figure S2:**
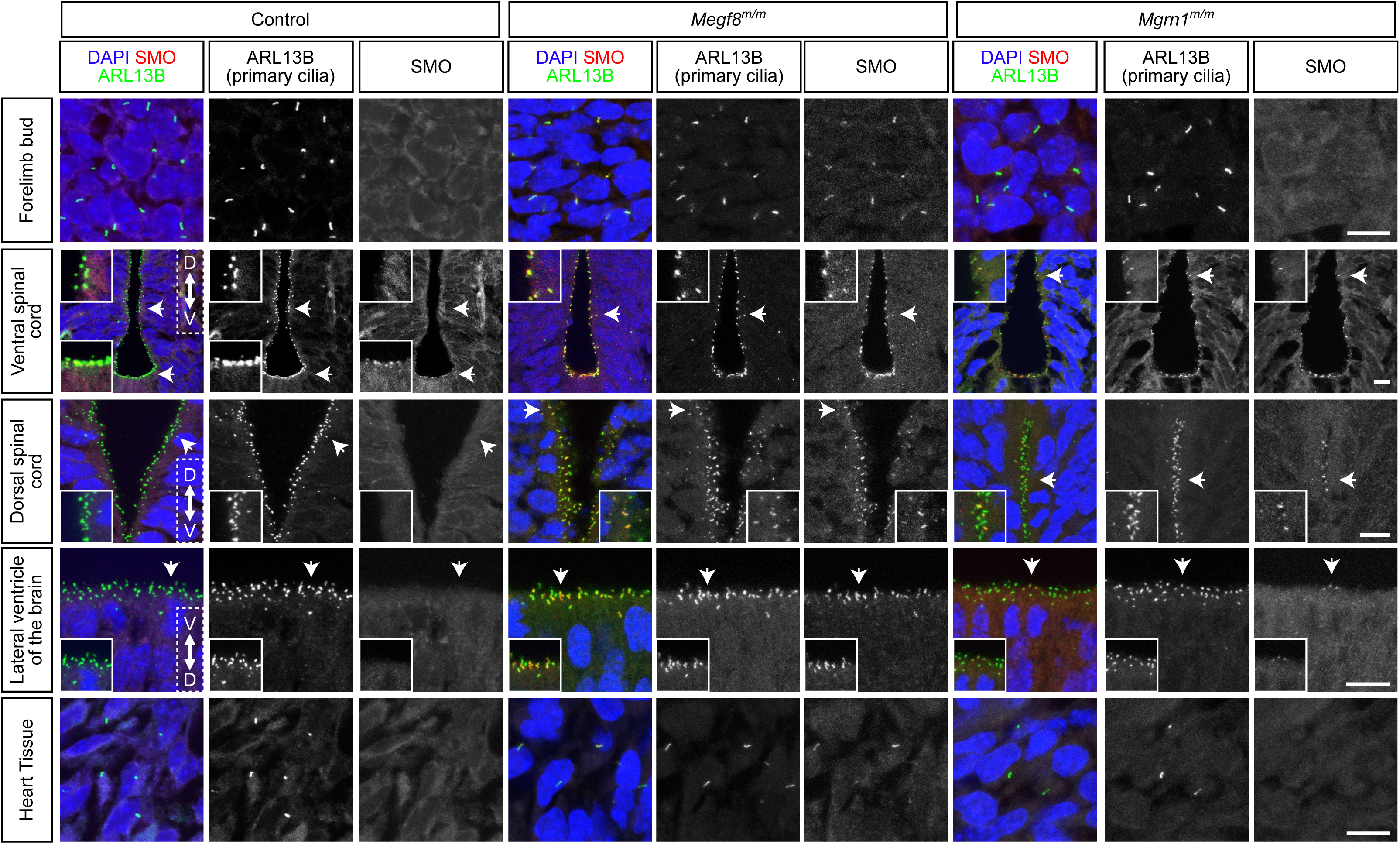
M*e*gf8m*^/m^* and *Mgrn1^m/m^* embryos have elevated ciliary Smoothened in multiple tissues, Related to Figure 1 Confocal fluorescence microscopy images of ciliated tissues (forelimb bud, spinal cord, lateral ventricles of the brain, and heart) collected from e12.5 control (*Mgrn1^m/+^* or *Megf8^m/+^*), *Megf8^m/m^*, and *Mgrn1^m/m^* embryos. Arrows denote regions enlarged in insets. In control embryos, ciliary SMO was only present in the floor plate of the ventral spinal cord. In contrast, SMO was detected at cilia in all *Megf8^m/m^* tissues analyzed. SMO was sparsely detected at cilia in the *Mgrn1^m/m^* spinal cord and brain. Red, SMO; green, ARL13B (primary cilia); blue, DAPI (nuclei). Scale bars, 10 µm.

**Figure S3:**
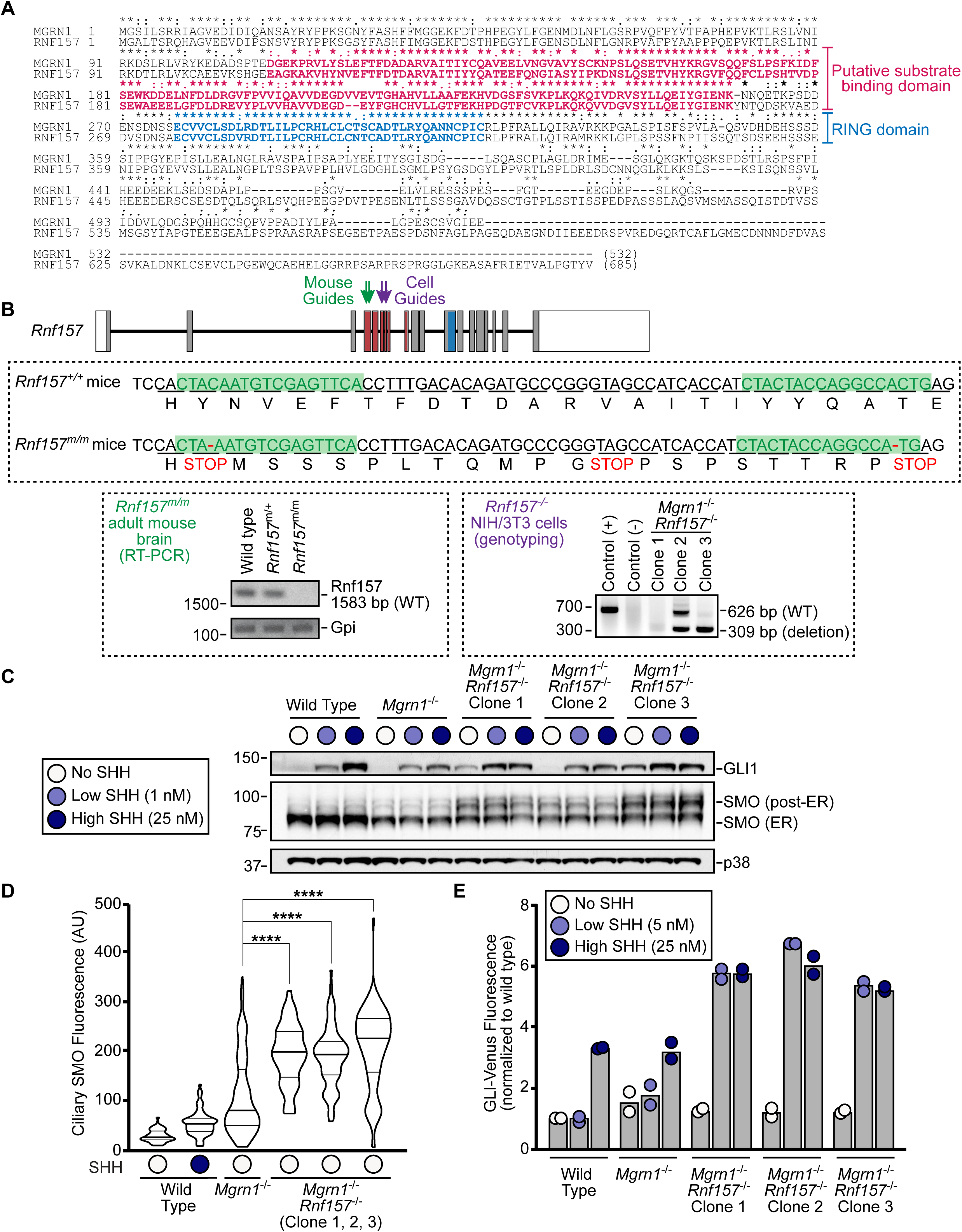
Loss-of-function mutations in both *Mgrn1* and *Rnf157* results in elevated Hh signaling strength in NIH/3T3 fibroblasts and neural progenitor cells, Related to Figure 1 **(A)** Alignment of mouse MGRN1 and RNF157. Red highlights a putative substrate-binding domain (Pusapati et al., 2018a) unique to proteins related to MGRN1 (tree shown in Fig. 1B), but not found in other RING superfamily E3 ligases. Blue highlights the RING domain. * fully conserved;: strongly similar (scoring > 0.5 in the Gonnet PAM 250 matrix);. weakly similar (scoring ≤ 0.5 in the Gonnet PAM 250 matrix). **(B)** CRISPR strategies for targeting *Rnf157* in mice (left, green) and in NIH/3T3 and neural progenitor cells (NPCs) (right, purple). Arrows denote the locations in the genomic map of *Rnf157* targeted by sgRNA guides in mice (green) and cells (purple). Open rectangles denote non-coding exons, gray rectangles denote coding exons, red rectangles denote regions that encode the putative substrate binding domain, blue rectangles denote regions that encode the RING domain, and horizontal lines denote introns. For the generation of *Rnf157^-/-^* mice, Sanger sequencing was used to identify a 2bp deletion that results in a premature stop codon (sequences are provided in the inset below the exon-intron map). RT-PCR was performed to validate the loss of *Rnf157* expression (bottom left). For the validation of *Rnf157^-/-^* NIH/3T3 and NPCs, PCR was used to confirm the deletion introduced by co-transfection of two sgRNA guides into the cells (bottom right). **(C and D)** Analysis of *Mgrn1*^-/-^;*Rnf157*^-/-^ NIH/3T3 cells. (C) Immunoblots showing the abundance of Hh signaling components in extracts of wild type, *Mgrn1^-/-^*, and three clonal *Mgrn1*^-/-^;*Rnf157*^-/-^ NIH/3T3 cell lines treated with no, low (1 nM), or high (25 nM) concentrations of SHH. (D) Violin plots summarize SMO fluorescence in ∼40 cilia per sample. Statistical significance was determined by the Kruskal-Wallis test; \*\*\*\**p*-value ≤ 0.0001. Clone 3 is featured in the main figures and was used to generate the stable cell lines shown in Fig. 2. **(E)** Hh signaling was assessed in Neural Precursor Cell lines (NPCs) of the indicated genotype exposed to no, low (5 nM), or high (25 nM) concentrations of SHH using a stably integrated fluorescent reporter of target gene induction (GLI-Venus, see (Pusapati et al., 2018a)). Bars represent the median GLI-Venus fluorescence (collected from 20,000 cells) from two independent experiments.

**Figure S4:**
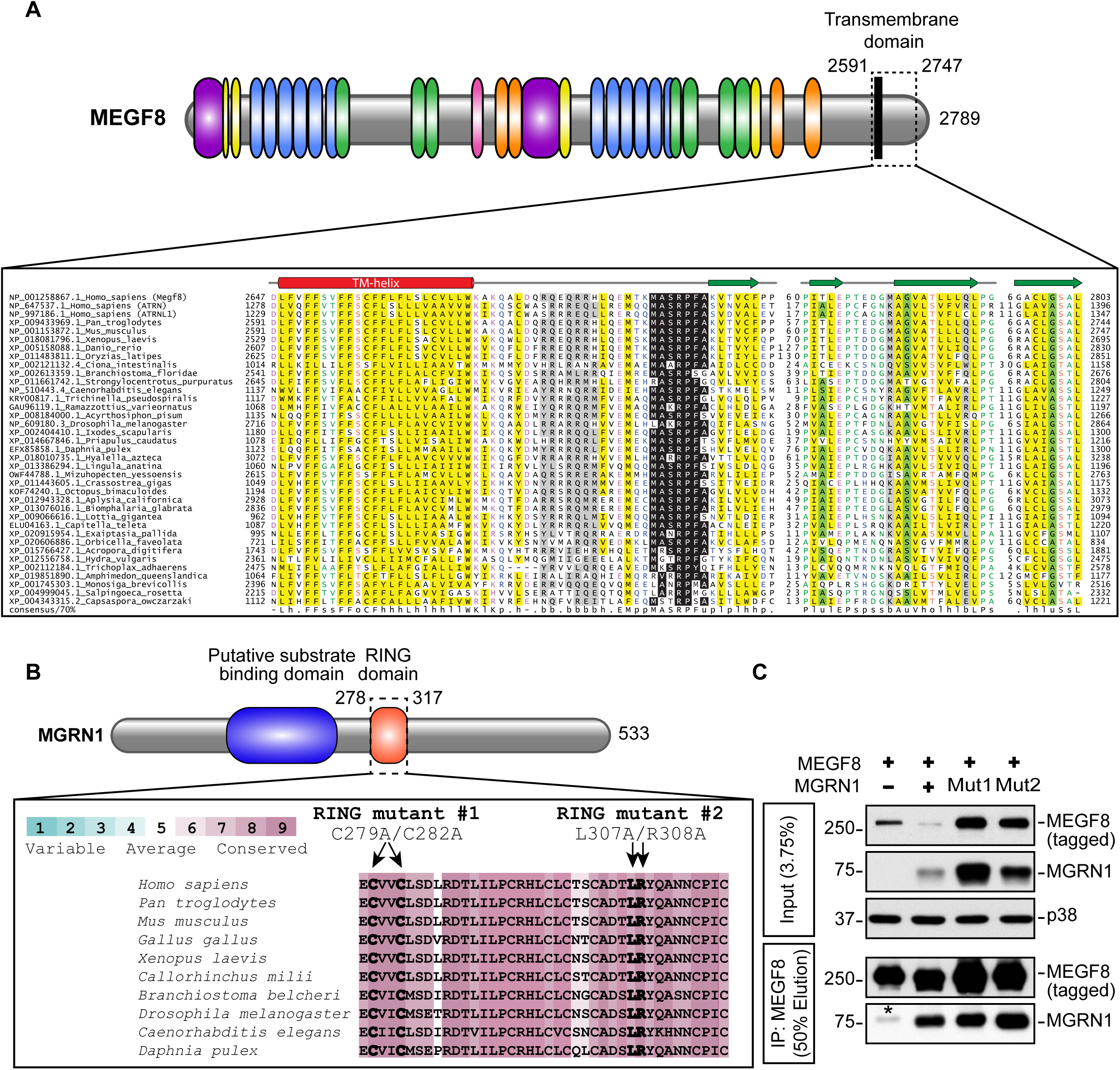
Sequence conservation in selected domains of MGRN1 and MEGF8, Related to Figure 2 **(A)** Sequence alignment showing conservation and secondary structure features of the cytoplasmic tail of several members of the MEGF8-ATRN family. The top three proteins are three paralogs (MEGF8, Attractin (ATRN) or Attractin-like (ATRNL1) found in humans and the rest are homologs of these proteins identified in various species. Proteins are named by their GenBank identifiers (GIs), followed by the complete names of the species from which they are derived. Predicted secondary structure elements are shown at the top: the red rectangle denotes a predicted α-helix and the multiple green arrows denote predicted β-strands. Low complexity regions between the strands are hidden and denoted as gaps with numbers. The MASRPFA sequence motif is highlighted in black. The alignment is colored based on 70% consensus with the following scheme as shown in the consensus sequence: h (hydrophobic), l (aliphatic), and a (aromatic) are shaded yellow; p (polar) are shaded blue; charged are shaded pale violet; s (small) and u (tiny) are shaded light green; b (big) is shaded dark gray. **(B)** ClustalW alignment of the RING domains of various MGRN1 homologs show conservation of the residues altered to generate the inactive MGRN1^Mut1^ (C279A/C282A) and MGRN1^Mut2^ (L307A/R308A) variants (Gunn et al., 2013b). Conservation at each position in the RING domain, color-coded according to the indicated scheme, is based on analysis of 200 homologs of MGRN1 using the ConSurf method (Ashkenazy et al., 2016). **(C)** Mutations in the RING domain introduced in MGRN1^Mut1^ and MGRN1^Mut2^ do not abolish their interactions with MEGF8 based on a co-IP experiment after transient expression of the indicated proteins in HEK293T cells (identical to the assay depicted in Fig. 2C). Asterisk (*) indicates endogenous MGRN1 from HEK293T cells that co-precipitated with MEGF8.

**Figure S5:**
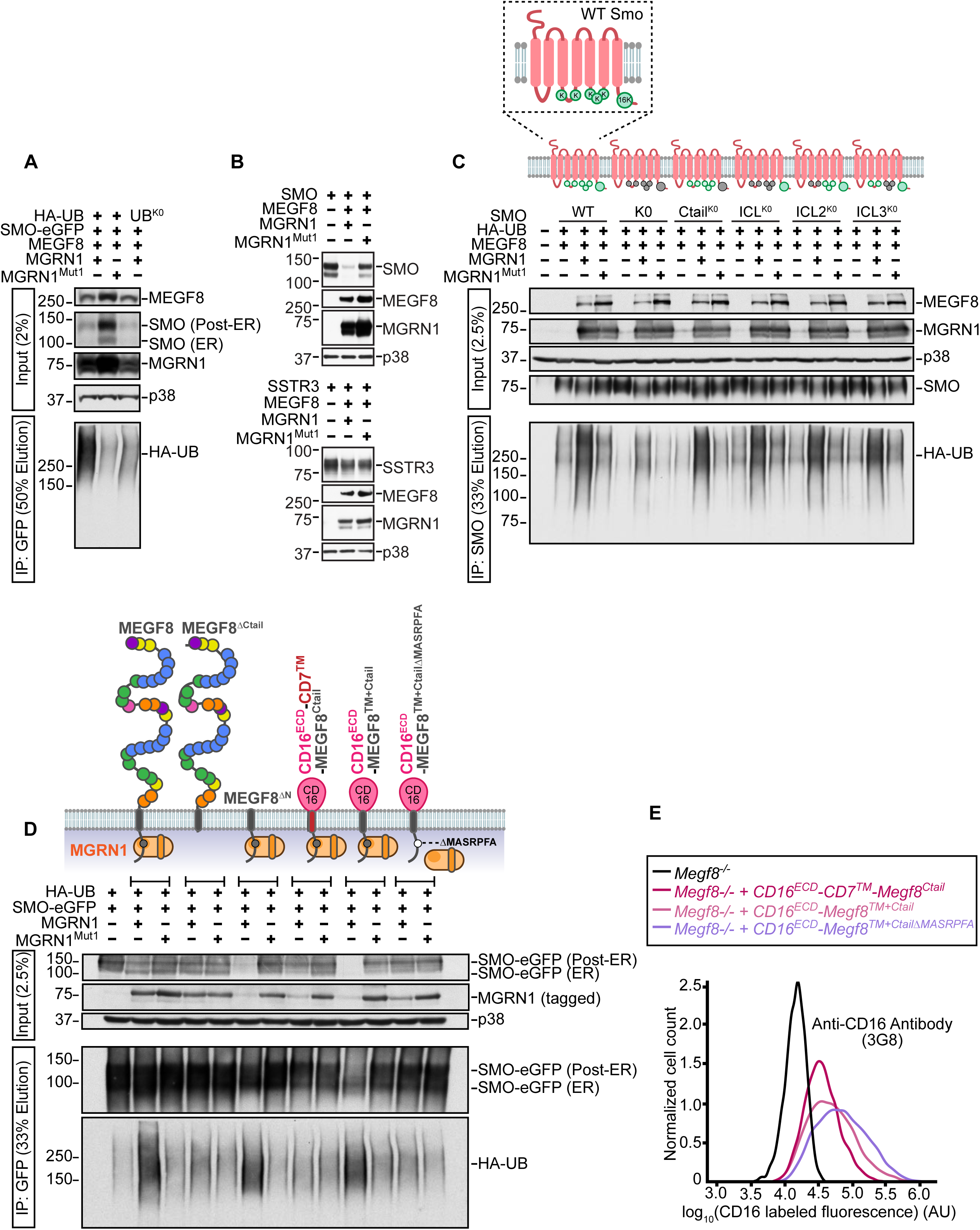
The MEGF8/MGRN1 complex ubiquitinates Smoothened, Related to Figure 4 **(A)** Ubiquitination of SMO-eGFP in the presence of wild-type HA-Ubiquitin (HA-UB) or a mutant where all the lysine residues have been mutated to arginine (UB^K0^) to prevent ubiquitin chain elongation. Ubiquitination was assessed as described in Fig. 4A after transient expression of the indicated proteins in HEK293T cells. **(B)** To determine if the MEGF8-MGRN1 complex can regulate the abundance of other G-protein coupled receptors (GPCRs), HEK293T cells were transiently transfected with constructs encoding MEGF8, functional or catalytically inactive MGRN1 (MGRN1^Mut1^), and either SMO (top) or Somatostatin receptor type 3 (SSTR3) (bottom). Immunoblots indicate that co-expression of MEGF8 and MGRN1 complex reduced the abundance of SMO, but had no effect on SSTR3. **(C)** Ubiquitination of wild type SMO or variants carrying mutations in cytoplasmically-exposed lysine residues by the MEGF8-MGRN1 complex. The following five SMO mutants were tested: (1) SMO-K0 (all 21 cytoplasmic lysine residues were changed to arginines), (2) Smo-Ctail^K0^ (all 16 lysines in the C-terminal cytoplasmic tail were changed to arginines), (3) SMO-ICL^K0^ (all 5 lysines in the three cytoplasmic loops were changed to arginine), (4) SMO-ICL2^K0^ (both lysines in the second intracellular loop were changed to arginines), and (5) SMO-ICL3^K0^ (all three lysines in the third intracellular loop were changed to arginines). Cells were lysed under denaturing conditions, native SMO was purified by IP using beads covalently linked to an anti-SMO antibody, and the amount of HA-UB covalently conjugated to SMO was assessed using immunoblotting with an anti-HA antibody (bottom panel). **(D)** Chimeric proteins were used to identify the minimal region of MEGF8 sufficient to support SMO ubiquitination. Using the assay shown in Fig. 4A, SMO ubiquitination was assessed after transient co-expression of the following in 293T cells: SMO-eGFP, HA-UB, MGRN1 (or the inactive mutant MGRN1^Mut1^), and the MEGF8 mutant or chimera shown above the blot. See Fig. 2A and associated text for a description of these chimeras. **(E)** Flow cytometry was used to measure cell surface labeled CD16 in live *Megf8^-/-^* cells stably expressing various CD16/CD7/MEGF8 chimeras (diagrammed in **Fig. S5D**). These are the same stable cell lines analyzed in Figs. **4C and 4D**. 4000 cells were analyzed per condition.

**Figure S6:**
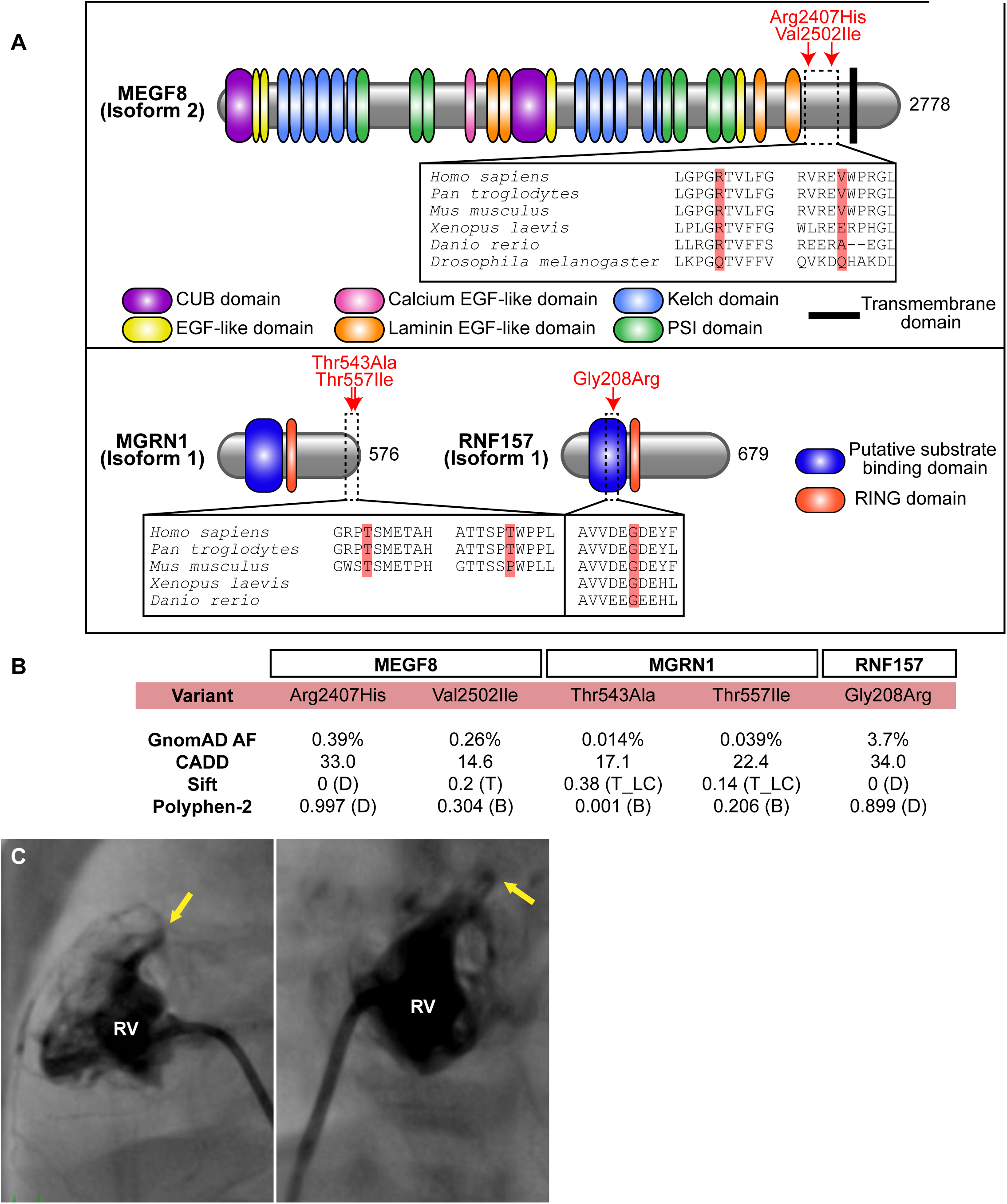
Analysis of human variants in *MEGF8*, *MGRN1*, and *RNF157*, Related to Figure 7 **(A)** Schematic representation of the domain composition of human MEGF8, MGRN1, and RNF157 (UniProt Consortium, 2019). Red arrows denote the location of the variants found in patient 7501 (described in Fig. 7A). The conservation of the amino acid residues around the variant of interest is analyzed across model species, with the specific residue altered by the variation shaded in red. **(B)** Population allele frequency (AF) of the variants found in patient 7501 is shown from the Genome Aggregation Database (GnomAD). The predicted deleteriousness of each variation on protein structure and function is estimated based on scores from three algorithms: Combined Annotation Dependent Depletion (CADD), Polymorphism Phenotyping v2 (Polyphen-2), or Sorting Intolerant from Tolerant (SIFT). B, benign; D, deleterious; T, tolerated; T_LC, is tolerated (low confidence). **(C)** Ventriculogram from patient 7501 in postero-anterior (left) and lateral (right) views demonstrated severely hypoplastic and hypertrophied right ventricle (RV) with an diminutive right ventricular outflow tract. There are copious right ventricular sinusoids coronary fistulas with what is seen of the coronary circulation returning to the aortic root. There was no antegrade flow from the right ventricle to the main pulmonary artery (yellow arrow).

**Table S1:** Heart and Visceral Organ Situs of Mgrn1^m/m^, Rnf157^m/m^, Megf8^m/m^, and Mgrn1^m/m^;Rnf157^m/m^ mice, Related to Figure 1

| Embryo ID | Body | Heart | VA Alignment | Septal Defect | Aortic Arch | Lung Situs | Abdomen Situs | RF | LF | RH | LH |
| --- | --- | --- | --- | --- | --- | --- | --- | --- | --- | --- | --- |
| 1 | SS | Lev | CC | - | LAA | 4R:1L | L(St/Sp/P) | NML | NML | NML | NML |
| 2 | SS | Lev | CC | - | LAA | 4R:1L | L(St/Sp/P) | NML | NML | NML | NML |
| 3 | SS | Lev | CC | - | LAA | 4R:1L | L(St/Sp/P) | NML | NML | NML | NML |
| 4 | SS | Lev | CC | - | LAA | 4R:1L | L(St/Sp/P) | NML | NML | NML | NML |
| 5 | SS | Lev | CC | - | LAA | 4R:1L | L(St/Sp/P) | NML | NML | NML | NML |
| 6 | SS | Lev | CC | - | LAA | 4R:1L | L(St/Sp/P) | NML | NML | NML | NML |

***Mgrn1<sup>m/m</sup>* mice**
| Embryo ID | Body | Heart | VA Alignment | Septal Defect | Aortic Arch | Lung Situs | Abdomen Situs | RF | LF | RH | LH |
| --- | --- | --- | --- | --- | --- | --- | --- | --- | --- | --- | --- |
| 1 | SIT | Dex | CC | VSD(Ao) | RAA | 1R:4L:SIT | R(St/Sp/P) | PDD | PDD | PDD | PDD |
| 2 | SS | Lev | CC | - | LAA | 4R:1L | L(St/Sp/P) | NML | NML | NML | NML |
| 3 | SS | Lev | CC | - | LAA | 4R:1L | L(St/Sp/P) | NML | NML | NML | NML |
| 4 | SS | Lev | CC | - | LAA | 4R:1L | L(St/Sp/P) | NML | NML | NML | NML |
| 5 | HTX | Dex | TGA | AVSD | RAA | 3R:3L:1M:RPI | L(St/Sp/P) | NML | NML | NML | NML |
| 6 | HTX | Dex | TGA | AVSD | RAA | 1R:1L:LPI | L(St/Sp/P) | NML | NML | NML | NML |
| 7 | SS | Lev | CC | - | LAA | 4R:1L | L(St/Sp/P) | NML | NML | NML | NML |
| 8 | SS | Lev | CC | - | LAA | 4R:1L | L(St/Sp/P) | NML | NML | NML | NML |
| 9 | SS | Lev | CC | - | LAA | 4R:1L | L(St/Sp/P) | NML | NML | NML | NML |
| 10 | SIT | Dex | TGA | AVSD | RAA | 1R:4L:SIT | R(St/Sp/P) | NML | NML | NML | NML |
| 11 | SS | Lev | CC | - | LAA | 4R:1L | L(St/Sp/P) | NML | NML | NML | NML |
| 12 | SS | Lev | CC | - | LAA | 4R:1L | L(St/Sp/P) | NML | NML | NML | NML |
| 13 | SS | Lev | CC | - | LAA | 4R:1L | L(St/Sp/P) | NML | NML | NML | NML |
| 14 | SS | Lev | CC | - | LAA | 4R:1L | L(St/Sp/P) | NML | NML | NML | NML |
| 15 | SS | Lev | CC | - | LAA | 4R:1L | L(St/Sp/P) | NML | NML | NML | NML |

***Megf8<sup>m/m</sup>* mice**
| Embryo ID | Body | Heart | VA Alignment | Septal Defect | Aortic Arch | Lung Situs | Abdomen Situs | RF | LF | RH | LH |
| --- | --- | --- | --- | --- | --- | --- | --- | --- | --- | --- | --- |
| 1 | HTX | Dex | TGA | AVSD | RAA | 3R:3L:1M:RPI | L(St/Sp/P) | PDD | PDD | PDD | PDD |
| 2 | HTX | Mes | TGA | AVSD | RAA | 3R:3L:1M:RPI | R(St/Sp/P) | PDD | PDD | PDD | PDD |
| 3 | HTX | Mes | TGA | VSD(Ao) | LAA | 3R:3L:1M:RPI | R(St/Sp/P) | N/A | PDD | PDD | PDD |
| 4 | HTX | Dex | TGA | AVSD | RAA | 1R:1L:LPI | L(St/Sp/P) | PDD | PDD | N/A | PDD |
| 5 | HTX | Lev | TGA | AVSD | LAA | 3R:3L:1M:RPI | R(St/Sp/P) | PDD | PDD | PDD | PDD |
| 6 | HTX | Dex | TGA | - | RAA | 3R:3L:1M:RPI | R(St/Sp/P) | PDD | PDD | N/A | PDD |
| 7 | HTX | Mes | TGA | AVSD | LAA | 3R:3L:1M:RPI | R(St/Sp/P) | PDD | PDD | PDD | PDD |
| 8 | HTX | Lev | TGA | AVSD | LAA | 3R:3L:1M:RPI | R(St/Sp/P) | PDD | N/A | PDD | PDD |
| 9 | HTX | Dex | TGA | - | RAA | 3R:3L:1M:RPI | L(St/Sp/P) | PDD | PDD | PDD | PDD |
| 10 | HTX | Lev | TGA | AVSD | LAA | 3R:3L:1M:RPI | R(St/Sp/P) | PDD | PDD | PDD | PDD |
| 11 | HTX | Lev | TGA | AVSD | LAA | 4R:1L | R(St/Sp/P) | PDD | PDD | PDD | PDD |
| 12 | HTX | Dex | TGA | AVSD | RAA | 3R:3L:1M:RPI | L(St/Sp/P) | PDD | PDD | PDD | PDD |

***Mgrn1<sup>m/m</sup>;Rnf157<sup>m/m</sup>* double knockout mice**
| Embryo ID | Body | Heart | VA Alignment | Septal Defect | Aortic Arch | Lung Situs | Abdomen Situs | RF | LF | RH | LH |
| --- | --- | --- | --- | --- | --- | --- | --- | --- | --- | --- | --- |
| 1 | HTX | Lev | TGA | AVSD | LAA | 3R:3L:1M:RPI | L(St/Sp/P) | PDD | PDD | PDD | PDD |
| 2 | HTX | Dex | TGA | AVSD | RAA | 3R:3L:1M:RPI | L(St/Sp/P) | N/A | PDD | NML | NML |
| 3 | HTX | Lev | TGA | AVSD | RAA | 3R:3L:1M:RPI | L(St/Sp/P) | PDD | PDD | PDD | PDD |

AVSD, atrioventricular septal defect; CC, concordant ventriculoarterial alignment; Dex, dextrocardia; DORV, double outlet right ventricle; HAA, hypoplastic arch; HTA, hypoplastic transverse heart; HTX, heterotaxy; IAA, interrupted aortic arch; LAA, left aortic arch; Lev, levocardia; LF, left forelimb; LH, left hindlimb; LPI, left pulmonary isomerism; L(St/Sp/P), Left sided stomach, spleen, pancreas; Mes, mesocardia; N/A: no digit data collected; NML, normal digit pattern; PDD, preaxial digit duplication; RAA, right aortic arch; RF, right forelimb; RH, right hindlimb; RPI, right pulmonary isomerism; R(St/Sp/P), Right sided stomach, spleen, pancreas; SIT, situs inversus; SS, situs solitus; SymLiv, symmetric liver; TGA, transposition of the great arteries; VA, ventriculoarterial; VSD, ventricular septal defect; VSD(Ao), ventricular septal defect located below the aorta; 1L, one lung lobe on left side; 4R: 4 lung lobes on right side.

**Table S2:** Heart and Visceral Organ Situs in Megf8^m/+^ single heterozygous mice, Related to Figures 5 and 6

| Embryo ID | Body | Heart | VA Alignment | Septal Defect | Aortic Arch | Lung Situs | Abdomen Situs | RF | LF | RH | LH |
| --- | --- | --- | --- | --- | --- | --- | --- | --- | --- | --- | --- |
| 1 | SS | Lev | CC | - | LAA | 4R:1L | L(St/Sp/P) | NML | NML | NML | N/A |
| 2 | SS | Lev | CC | - | LAA | 4R:1L | L(St/Sp/P) | N/A | NML | NML | N/A |
| 3 | SS | Lev | CC | - | LAA | 4R:1L | L(St/Sp/P) | NML | NML | NML | NML |
| 4 | SS | Lev | CC | - | LAA | 4R:1L | L(St/Sp/P) | N/A | N/A | NML | NML |
| 5 | SS | Lev | CC | - | LAA | 4R:1L | L(St/Sp/P) | NML | NML | NML | NML |
| 6 | SS | Lev | CC | - | LAA | 4R:1L | L(St/Sp/P) | NML | NML | NML | NML |
| 7 | SS | Lev | CC | - | LAA | 4R:1L | L(St/Sp/P) | NML | NML | N/A | N/A |
| 8 | SS | Lev | CC | - | LAA | 4R:1L | L(St/Sp/P) | NML | NML | N/A | NML |
| 9 | SS | Lev | CC | - | LAA | 4R:1L | L(St/Sp/P) | NML | NML | NML | NML |
| 10 | SS | Lev | CC | - | LAA | 4R:1L | L(St/Sp/P) | NML | NML | N/A | NML |
| 11 | SS | Lev | CC | - | LAA | 4R:1L | L(St/Sp/P) | NML | NML | NML | NML |
| 12 | SS | Lev | CC | - | LAA | 4R:1L | L(St/Sp/P) | NML | NML | NML | NML |
| 13 | SS | Lev | CC | - | LAA | 4R:1L | L(St/Sp/P) | NML | NML | NML | NML |
| 14 | SS | Lev | CC | - | LAA | 4R:1L | L(St/Sp/P) | N/A | NML | NML | NML |

CC, concordant ventriculoarterial alignment; LAA, left aortic arch; Lev, levocardia; LF, left forelimb; LH, left hindlimb; L(St/Sp/P), Left sided stomach spleen, pancreas; N/A, no digit data collected; NML, normal digit pattern; RF, right forelimb; RH, right hindlimb; SS, situs solitus, 1L, one lung lobe on left side; 4R, 4 lung lobes on right side.

**Table S3:** Heart and Visceral Organ Situs in *Mgrn1^m /+^* single heterozygous mice, Related to **Figures 5 and 6**

| Embryo ID | Body | Heart | VA Alignment | Septal Defect | Aortic Arch | Lung Situs | Abdomen Situs | RF | LF | RH | LH |
| --- | --- | --- | --- | --- | --- | --- | --- | --- | --- | --- | --- |
| 1 | SS | Lev | CC | - | LAA | 4R:1L | L(St/Sp/P) | N/A | NML | NML | N/A |
| 2 | SS | Lev | CC | - | LAA | 4R:1L | L(St/Sp/P) | NML | NML | NML | NML |
| 3 | SS | Lev | CC | - | LAA | 4R:1L | L(St/Sp/P) | NML | NML | NML | NML |
| 4 | SS | Lev | CC | - | LAA | 4R:1L | L(St/Sp/P) | NML | NML | N/A | N/A |
| 5 | SS | Lev | CC | - | LAA | 4R:1L | L(St/Sp/P) | N/A | N/A | NML | NML |
| 6 | SS | Lev | CC | - | LAA | 4R:1L | L(St/Sp/P) | NML | NML | NML | NML |
| 7 | SS | Lev | CC | - | LAA | 4R:1L | L(St/Sp/P) | NML | NML | NML | NML |
| 8 | SS | Lev | CC | - | LAA | 4R:1L | L(St/Sp/P) | N/A | N/A | N/A | NML |
| 9 | SS | Lev | CC | - | LAA | 4R:1L | L(St/Sp/P) | NML | NML | NML | NML |
| 10 | SS | Lev | CC | - | LAA | 4R:1L | L(St/Sp/P) | NML | NML | NML | NML |
| 11 | SS | Lev | CC | - | LAA | 4R:1L | L(St/Sp/P) | NML | NML | NML | NML |
| 12 | SS | Lev | CC | - | LAA | 4R:1L | L(St/Sp/P) | N/A | NML | NML | NML |
| 13 | SS | Lev | CC | - | LAA | 4R:1L | L(St/Sp/P) | NML | NML | N/A | NML |
| 14 | SS | Lev | CC | - | LAA | 4R:1L | L(St/Sp/P) | NML | NML | N/A | N/A |
| 15 | SS | Lev | CC | - | LAA | 4R:1L | L(St/Sp/P) | NML | NML | NML | NML |

CC, concordant ventriculoarterial alignment; LAA, left aortic arch; Lev, levocardia; LF, left forelimb; LH, left hindlimb; L(St/Sp/P), Left sided stomach spleen, pancreas; N/A, no digit data collected; NML, normal digit pattern; RF, right forelimb; RH, right hindlimb; SS, situs solitus, 1L, one lung lobe on left side; 4R, 4 lung lobes on right side.

**Table S4:** Heart and Visceral Organ Situs in Megf8^m/+^;Mgrn1^m/+^ double heterozygous mice, Related to Figures 5 and 6

| Embryo ID | Body | Heart | VA Alignment | Septal Defect | Aortic Arch | Lung Situs | Abdomen Situs | RF | LF | RH | LH |
| --- | --- | --- | --- | --- | --- | --- | --- | --- | --- | --- | --- |
| 1 | SS | Lev | CC | - | LAA | 4R:1L | L(St/Sp/P) | PDD | PDD | N/A | N/A |
| 2 | SS | Lev | DORV | - | LAA | 4R:1L | L(St/Sp/P) | PDD | N/A | N/A | N/A |
| 3 | SS | Lev | CC | - | LAA | 4R:1L | L(St/Sp/P) | PDD | PDD | N/A | N/A |
| 4 | SS | Lev | CC | - | LAA | 4R:1L | L(St/Sp/P) | PDD | PDD | PDD | PDD |
| 5 | HTX | Dex | CC | - | LAA | 4R:1L | L(St/Sp/P) | NML | NML | NA | NML |
| 6 | HTX | Lev | DORV | VSD(Ao) | LAA,HAA, IAA | 1R:1L:LPI | L(St/Sp/P) | NML | NML | NML | NML |
| 7 | SS | Lev | CC | - | LAA | 4R:1L | L(St/Sp/P) | NML | NML | NML | NML |
| 8 | SS | Lev | CC | - | LAA | 4R:1L | L(St/Sp/P) | PD | PDD | N/A | N/A |
| 9 | SS | Lev | CC | - | LAA | 4R:1L | L(St/Sp/P) | NML | NML | NML | NML |
| 10 | SS | Lev | CC | AVSD | LAA | 2R:1L:LPI | L(St/Sp/P) | PDD | PDD | N/A | N/A |
| 11 | SS | Lev | CC | - | LAA | 4R:1L | L(St/Sp/P) | NML | NML | NML | NML |
| 12 | HTX | Lev | DORV | - | LAA | 1R:1L:LPI | L(St/Sp/P) | PDD | PD | N/A | N/A |
| 13 | SIT | Dex | DORV | VSD(Pa) | RAA | 4L:1R:SI | R(St/Sp/P) SymLiv | NML | NML | NML | NML |
| 14 | SS | Lev | CC | - | LAA | 4R:1L | L(St/Sp/P) | PDD | PDD | N/A | N/A |
| 15 | SS | Lev | DORV | AVSD | LAA | 4R:1L | L(St/Sp/P) | NML | NML | NML | NML |
| 16 | HTX | Lev | TGA | AVSD | LAA | 3R:3L:1M:RPI | R(St/Sp/P) SymLiv | NML | NML | PDD | NML |
| 17 | SS | Lev | CC | - | LAA | 4R:1L | L(St/Sp/P) | NML | NML | NML | NML |
| 18 | HTX | Lev | DORV | VSD(Pa) | LAA | 3R:3L:1M:RPI | L(St/Sp/P) | NML | NML | NML | NML |
| 19 | SS | Lev | DORV | VSD(Ao) | LAA,HTA | 4R:1L | L(St/Sp/P) | NML | NML | PDD | NML |
| 20 | HTX | Lev | TGA | VSD(Pa) | LAA | 3R:3L:1M:RPI | L(St/Sp/P) | PDD | PDD | PDD | PDD |
| 21 | HTX | Mal | TGA | AVSD | LAA | 3R:3L:1M:RPI | L(St/Sp/P) | PDD | PDD | N/A | N/A |
| 22 | HTX | Dex | TGA | AVSD | RAA | 3R:3L:1M:RPI | R(St/Sp/P) SymLiv | PDD | PDD | PDD | PDD |
| 23 | SS | Lev | CC | - | LAA | 4R:1L | L(St/Sp/P) | NML | NML | PDD | NML |
| 24 | SS | Lev | CC | - | LAA | 4R:1L | L(St/Sp/P) | NML | NML | NML | NML |
| 25 | SS | Lev | CC | - | LAA | 4R:1L | L(St/Sp/P) | NML | NML | NML | NML |
| 26 | SS | Lev | CC | - | LAA | 4R:1L | L(St/Sp/P) | PDD | PDD | PDD | PDD |
| 27 | SS | Lev | CC | - | LAA | 4R:1L | L(St/Sp/P) | NML | NML | NML | NML |
| 28 | HTX | Dex | TGA | AVSD | RAA | 3R:3L:1M:RPI | L(St/Sp/P) | PDD | PDD | PDD | PDD |
| 29 | SS | Lev | CC | - | LAA | 4R:1L | L(St/Sp/P) | NML | NML | NML | NML |
| 30 | HTX | Dex | TGA | AVSD | RAA | 3R:3L:1M:RPI | R(St/Sp/P) SymLiv | PDD | PDD | N/A | PDD |
| 31 | HTX | Lev | TGA | AVSD | LAA | 3R:3L:1M:RPI | R(St/Sp/P) SymLiv | PDD | N/A | PDD | PDD |
| 32 | SS | Lev | CC | - | LAA | 4R:1L | L(St/Sp/P) | PDD | PDD | PDD | PDD |
| 33 | SS | Lev | DORV | - | LAA | 4R:1L | L(St/Sp/P) | PDD | PDD | N/A | N/A |

AVSD, atrioventricular septal defect; CC, concordant ventriculoarterial alignment; Dex, dextrocardia; DORV, double outlet right ventricle; HAA, hypoplastic arch; HTA, hypoplastic transverse heart; HTX, heterotaxy; IAA, interrupted aortic arch; LAA, left aortic arch; Lev, levocardia; LF, left forelimb; LH, left hindlimb; LPI, left pulmonary isomerism; L(St/Sp/P), Left sided stomach, spleen, pancreas; Mal, malrotated; N/A: no digit data collected; NML, normal digit pattern; PDD, preaxial digit duplication; RAA, right aortic arch; RF, right forelimb; RH, right hindlimb; RPI, right pulmonary isomerism; R(St/Sp/P), Right sided stomach, spleen, pancreas; SIT, situs inversus; SS, situs solitus; SymLiv, symmetric liver; TGA, transposition of the great arteries; VA, ventriculoarterial; VSD, ventricular septal defect; VSD(Ao or Pa), ventricular septal defect located below the aorta or pulmonary artery; 1L, one lung lobe on left side; 4R: 4 lung lobes on right side.

**Table S5:** Heart and Visceral Organ Situs in *Megf8^m/+^;Mgrn1^m/m^* mice, Related to **Figure 6**

| Embryo ID | Body | Heart | VA Alignment | Septal Defect | Aortic Arch | Lung Situs | Abdomen Situs | RF | LF | RH | LH |
| --- | --- | --- | --- | --- | --- | --- | --- | --- | --- | --- | --- |
| 1 | HTX | Lev | DORV | AVSD | RAA | 3R:3L:1M:RPI | L(St/Sp/P) | N/A | PDD | PDD | PDD |
| 2 | HTX | Lev | DORV | VSD(Ao) | LAA | 3R:3L:1M:RPI | L(St/Sp/P) | PDD | PDD | PDD | PDD |
| 3 | SS | Lev | CC | - | LAA | 4R:1L | L(St/Sp/P) | PDD | PDD | PDD | PDD |
| 4 | SS | Lev | CC | - | LAA | 4R:1L | L(St/Sp/P) | NML | NML | NML | NML |
| 5 | HTX | Lev | TGA | AVSD | RAA | 3R:3L:1M:RPI | L(St/Sp/P) | PDD | PDD | PDD | PDD |
| 6 | SIT | Dex | TGA | AVSD | RAA | 1R:4L:SIT | R(St/Sp/P) | PDD | PDD | PDD | PDD |
| 7 | HTX | Dex | TGA | AVSD | RAA | 3R:3L:1M:RPI | L(St/Sp/P) | NML | NML | NML | NML |

AVSD, atrioventricular septal defect; CC, concordant ventriculoarterial alignment; Dex, dextrocardia; DORV, double outlet right ventricle; HTX, heterotaxy; LAA, left aortic arch; Lev, levocardia; LF, left forelimb; LH, left hindlimb; L(St/Sp/P), Left sided stomach, spleen, pancreas; N/A: no digit data collected; NML, normal digit pattern; PDD, preaxial digit duplication; RAA, right aortic arch; RF, right forelimb; RH, right hindlimb; RPI, right pulmonary isomerism; SIT, situs inversus; SS, situs solitus; TGA, transposition of the great arteries; VSD, ventricular septal defect; VSD(Ao), ventricular septal defect located below the aorta; 1L, one lung lobe on left side; 4R: 4 lung lobes on right side.

**Table S6:** Heart, limb, and laterality defects in *Megf8, Mgrn1,* and *Rnf157* mutant mice, Related to Figure 6

| Embryo genotype | CHD | DORV | TGA | AVSD | VSD | Dextrocardia | Heterotaxy | Situs inversus | Preaxial digit duplication |
| --- | --- | --- | --- | --- | --- | --- | --- | --- | --- |
| *Rnf157<sup>m/m</sup>*<br>n=6 | 0<br>(0%) | 0<br>(0%) | 0<br>(0%) | 0<br>(0%) | 0<br>(0%) | 0<br>(0%) | 0<br>(0%) | 0<br>(0%) | 0<br>(0%) |
| *Megf8<sup>m/+</sup>*<br>n=14 | 0<br>(0%) | 0<br>(0%) | 0<br>(0%) | 0<br>(0%) | 0<br>(0%) | 0<br>(0%) | 0<br>(0%) | 0<br>(0%) | 0<br>(0%) |
| *Mgrn1<sup>m/+</sup>*<br>n=15 | 0<br>(0%) | 0<br>(0%) | 0<br>(0%) | 0<br>(0%) | 0<br>(0%) | 0<br>(0%) | 0<br>(0%) | 0<br>(0%) | 0<br>(0%) |
| *Mgrn1<sup>m/m</sup>*<br>n=15 | 4<br>(27%) | 0<br>(0%) | 3<br>(20%) | 3<br>(20%) | 1<br>(7%) | 4<br>(27%) | 2<br>(13%) | 2<br>(13%) | 1<br>(7%) |
| *Megf8<sup>m/+</sup>*<br>*Mgrn1<sup>m/+</sup>*<br>n=33 | 17<br>(52%) | 8<br>(24%) | 7<br>(21%) | 8<br>(24%) | 5<br>(15%) | 5<br>(15%) | 11<br>(33%) | 1<br>(3%) | 20<br>(61%) |
| *Megf8<sup>m/+</sup>*<br>*Mgrn1<sup>m/m</sup>*<br>n=7 | 5<br>(71%) | 2<br>(29%) | 3<br>(43%) | 4<br>(57%) | 1<br>(14%) | 2<br>(29%) | 4<br>(57%) | 1<br>(14%) | 5<br>(71%) |
| *Megf8<sup>m/m</sup>*<br>n=12 | 12<br>(100%) | 0<br>(0%) | 12<br>(100%) | 9<br>(75%) | 1<br>(8%) | 5<br>(42%) | 12<br>(100%) | 0<br>(0%) | 12<br>(100%) |
| *Mgrn1<sup>m/m</sup>*<br>*Rnf157<sup>m/m</sup>*<br>n=3 | 3<br>(100%) | 0<br>(0%) | 3<br>(100%) | 3<br>(100%) | 0<br>(0%) | 1<br>(33%) | 3<br>(100%) | 0<br>(0%) | 3<br>(100%) |

AVSD, atrioventricular septal defect; DORV, double outlet right ventricle; CHD, congenital heart defect; TGA, transposition of the great arteries; VSD, ventricular septal defect

**Table S7:** Laterality of CHD lesions in *Megf8^m/+^;Mgrn1^m /+^* double heterozygous mice, Related to **Figure 6**

| **Specific CHD** | **Situs Solitus** | **Situs Inversus** | **Heterotaxy** |
| --- | --- | --- | --- |
| DORV<br>n=8 | 4<br>(50%) | 1<br>(12.5%) | 3<br>(37.5%) |
| TGA<br>n=7 | 0<br>(0%) | 0<br>(0%) | 7<br>(100%) |
| AVSD<br>n=8 | 2<br>(25%) | 0<br>(0%) | 6<br>(75%) |
| VSD<br>n=5 | 1<br>(20%) | 1<br>(20%) | 3<br>(60%) |

AVSD, atrioventricular septal defect; DORV, double outlet right ventricle; TGA, transposition of the great arteries; VSD, ventricular septal defect

**Table S8:** **Whole exome sequencing of patient 7501, Related to Figures 7**

| Sample ID | Gene | Chr:Position | Ref | Alt | Ref SNP | CADD PHREDD | GenomAD AF | Variant |
| --- | --- | --- | --- | --- | --- | --- | --- | --- |
| 7501 | MGRN1 | 16:4738805 | A | G | rs369718932 | 17.14 | 0.01377% | c.1627A>G:p.Thr543Ala |
| 7501 | MGRN1 | 16:4738848 | C | T | rs200375426 | 22.4 | 0.03913% | c.1670C>T:p.Thr557Ile |
| 7501 | RNF157 | 17:74162548 | C | T | rs11539879 | 34 | 3.695% | c.622G>A:p.Gly208Arg |
| 7501 | MEGF8 | 19:42879810 | G | A | rs45623135 | 33 | 0.3952% | c.7220G>A:p.Arg2407His |
| 7501 | MEGF8 | 19:42880094 | G | A | rs147133204 | 14.57 | 0.2581% | c.7504G>A:p.Val2502Ile |

